# New enhancer-promoter interactions are gained during tissue differentiation and reflect changes in E/P activity

**DOI:** 10.1101/2022.12.07.519443

**Authors:** Tim Pollex, Adam Rabinowitz, Maria Cristina Gambetta, Raquel Marco-Ferreres, Rebecca R. Viales, Aleksander Jankowski, Christoph Schaub, Eileen E.M. Furlong

## Abstract

To regulate gene expression, enhancers must come into proximity with their target gene. At some loci the timing of enhancer-promoter proximity is uncoupled from gene activation, while at others it is tightly linked. Here, we assessed this more globally for 600 characterized enhancers or promoters (E/P) with tissue-specific activity in *Drosophila* embryos, by performing Capture-C and insulator ChIP in FACS-purified myogenic or neurogenic cells at different stages of embryogenesis. This high-resolution view enabled direct comparison between E/P interactions and activity across 5 developmental conditions. This revealed largely invariant E/P contacts between the blastoderm and cell fate specification stages, despite changes in activity. However, E/P interactions diverge during terminal tissue differentiation when many tissue-specific interactions are gained on top of a pre-existing topology. Changes in E/P proximity reflect changes in enhancer activity and gene activation, and are generally not accompanied by changes in insulator binding. Using transgenes and deletions, we show that many tissue-specific interactions represent functional E-P pairs. Our results reveal a shift in E-P landscapes as embryogenesis proceeds, from largely pre-formed topologies at early stages to more distal tissue-specific loops during differentiation, when E/P proximity appears coupled to activation.

## INTRODUCTION

How distal enhancers convey regulatory information across large linear genomic distances to their target gene’s promoter has been intensely studied during the past decade. The prevailing model involves bringing the enhancer and promoter into spatial proximity through the formation of enhancer-promoter (E-P) loops. How and when these looping interactions are formed, and their relationship to gene activation remains unclear, especially within the context of embryonic development.

At some loci, enhancers appear to be only in proximity (or looped) to their target gene’s promoter in the appropriate cell type or developmental stage, where the gene is expressed^1–3^. For example, when comparing mouse embryonic stem cells with *in vitro* differentiated, or *in vivo* isolated, neural progenitor cells or neurons, many E-P interactions were more frequent at the stage when the gene was expressed, and the interaction strength correlated with the level of the gene’s expression^3^. Cell- and stage-specific chromatin interactions have also been observed during cardiac development^2^, adipocyte differentiation^4,5^ and at rhythmically expressed gene loci^6^. This suggests that the timing of E-P loop formation can be instructive for gene expression. In line with this, forcing an E-P interaction by tethering an enhancer and promoter together can even be sufficient to activate gene expression in both vertebrates^7–11^ and *Drosophila*^12^.

However, this coupling between the timing of E-P loop formation and the initiation of the gene’s expression is not observed at other loci or in other contexts, indicating that the formation of E-P interactions can be temporally separated from transcriptional activation.

Comparing the proximity of embryonic enhancers that are active during cell fate specification to the earlier blastoderm stage of *Drosophila* embryogenesis, in tightly staged 2-hour windows, revealed that the vast majority of enhancers were already in proximity to their target promoter hours before the gene is expressed^13^. Such constitutive or pre-formed E-P loops have also been observed in many other contexts including Zebrafish development^14^, mammalian cellular differentiation^15^ and limb development^16^, and in cell culture models during *trans*-differentiation^17^ and iPSC reprogramming^18^. In early *Drosophila* embryos, these pre-formed loops involved promoters with paused RNA polymerase II, suggesting that they are poised at the blastoderm stage, ready for rapid activation during the subsequent cell fate specification stages^13^. In line with this, pre-formed E-P loops have also been observed in the context of inducible gene expression^19,20^. Activity-dependent activation of gene expression in neurons is also associated with only a small fraction (∼10%) of *de novo* promoter loops, the majority do not change^21^.

This suggests that both types of E-P topologies, either tissue-specific or constitutive (appearing invariant between cell types or time-points), can exert cell-type specific responses^22,23^. For example, examining promoter interactions across different blood cell types^24^ or in response to diet adaptation^25^ revealed both types of interactions. This suggests that pre-formed topologies are perhaps established by ubiquitously expressed TFs, as suggested for CTCF/cohesin in the limb bud^16^, while the more dynamic instructive loops could be initiated by cell type specific TFs, (e.g. the neuronal pioneer factor Neurog2^26^), although few such factors have been identified or functionally tested to date. This is also likely an oversimplification, as many pre-formed loops are also associated with cell type-specific TFs, as seen during the differentiation of primary human keratinocytes^27^. However, there is currently little information about E-P topologies at later stages of a tissue’s terminal differentiation *in vivo*, within the context of embryonic development.

Here, we directly assessed this by examining the dynamic changes in E-P interactions during the transition from cell fate specification to tissue differentiation, during 2-hour windows of *Drosophila* embryonic development. We focused on a hand-curated set of ∼300 enhancers and ∼300 promoters with tissue-specific activity in either the muscle or nervous system at different stages. Capture-C was performed on this collection of regulatory elements (baits) in nuclei from myoblasts/muscle or neurogenic tissue/neurons from tightly staged embryos purified using fluorescence activated nuclei sorting (FANS, as described in ^28^). We then quantified the temporal dynamics and tissue-specificity of E-P interactions during three critical stages of embryonic development: early blastoderm (when cells are multipotent), cell fate specification of myoblasts or neurogenic system, and terminal differentiation to muscles and neurons. This high-resolution view of E/P interactions across tissues and stages revealed that there are few changes between the blastoderm and cell fate specification stages, with the majority of E/P interactions being already present in the blastoderm embryo, as we reported previously^13^. Moreover, specified myoblasts and neurons have surprisingly similar enhancer-promoter topologies between each other (at 6-8h), even though they express cell-type specific identity markers. However, at later stages of embryogenesis, during terminal tissue differentiation of the muscle and neurons, regulatory landscapes become more diverged and tissue-specific, where new E-P loops emerge on-top of the pre-formed topologies. Formation of these *de novo* interactions is associated with a gain in activity, with concordant changes in E/P interaction frequencies and the regulatory elements activity and chromatin state, suggesting that are functional regulatory events. We show using transgenic reporter assays that many differential loops with active promoters function as developmental enhancers *in vivo* in the appropriate tissues and stages. Deletion of two of these elements caused developmental defects, particularly under environmental stress.

Taken together, our results indicate that while the transition from pluripotency to cell fate specification involves pre-formed, largely invariant, E-P topologies, the transition from specified cell types to differentiated tissues involves many *de novo*, often more distal E-P contacts that are formed on-top of pre-existing topologies. At differentiation stages, these changes in E/P proximity generally reflect changes in enhancer activity and gene activation. Preformed E/P topologies may enable rapid development at early embryonic stages, while at later stages the use of more distal tissue-specific elements might give more robustness and stability to developmental programs.

## RESULTS

### Quantifying enhancer and promoter interactions across different tissues and stages of embryogenesis

To dissect the relationship between enhancer activity, enhancer-promoter (E-P) proximity and gene expression we performed high-resolution Capture-C focusing on over 600 characterised developmental enhancers or promoters with tissue- and stage-specific activity (Fig. 1a). We selected enhancers and promoters active during myoblast specification (Myo, including somatic and visceral muscle) or differentiated muscle tissues, or in the developing embryonic nervous system (neuro) or differentiated neurons (Fig. 1b), and assessed their 3D genome organization (as measured by cross-linking frequency in Capture-C) in five different developmental contexts (Fig. 1a). Specifically, Capture-C was performed on nuclei isolated from (1) early blastoderm stage embryos, a timepoint when cells are multipotent (mainly stage 5), (2) from mid-stage embryos during the specification of major lineages (6-8h after egg laying, mainly stage 10/11) and (3) later-stage embryos during the initiation of terminal tissue differentiation (10-12h after egg laying, mainly stage 13). The isolated nuclei from the latter two time-points were stained with antibodies specific for a nuclear marker of myoblast and muscle cells in the developing mesoderm (Mef2) and cells in the developing nervous system (Elav), and FAN sorted to over 95% purity using our optimised BiTS protocol^28^, allowing us to perform tissue-and stage-specific Capture-C (Fig. 1a, Methods).

**Figure 1:**
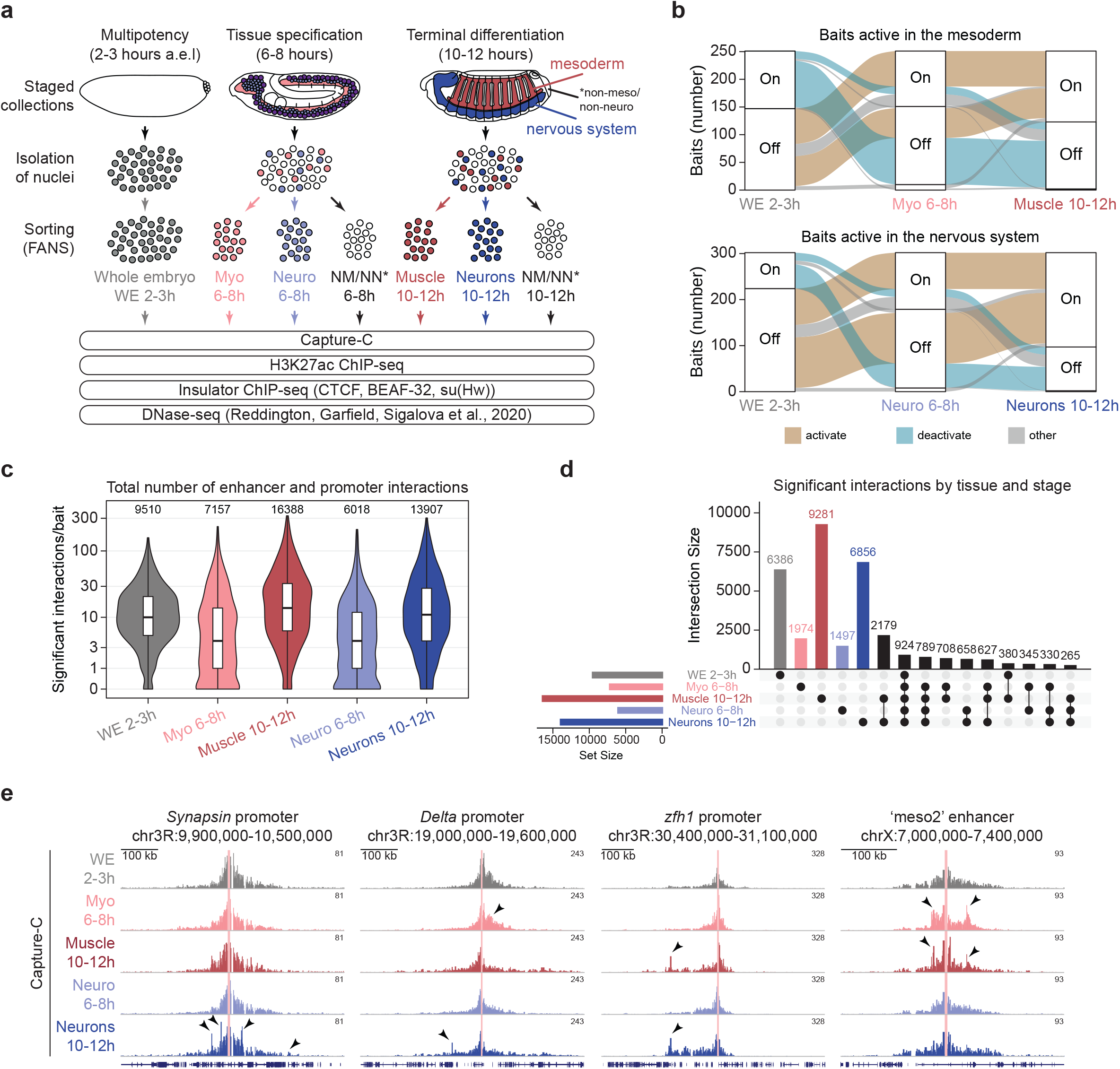
Quantifying enhancer and promoter interactions across tissues and developmental time. **(A)** Experimental overview: Myogenic (Mef2+) and Neurogenic (Elav+) cells were isolated (>95% purity) from tightly staged embryos and used for Capture-C and ChIP-seq for H3K27ac, CTCF, BEAF-32 and Su(Hw) in biological replicates. Grey = whole embryo 2-3h (WE 2-3h), red = myogenic mesoderm (Myo 6-8h, Muscle 10-12h), blue = nervous system (Neuro 6-8h, Neurons 10-12h). NM/NN = non-meso non-neuro (Mef2- & Elav-cells). Alluvial plots showing the dynamic activity of the enhancer/promoter baits in muscle and/or neuronal systems (some are dynamic in both tissues). **(C)** Violin plots of the number of significant interactions (CHiCAGO score >5) per bait at the indicated developmental time/tissue (colors as in (A)). Total number of significant interactions for all baits indicated above. **(D)** UpSet plot showing significant enhancer or promoter interactions (CHiCAGO score >5) in the five tested tissue- and stage-conditions. Unique interactions for each condition are indicated by colored bars (as in (A). Only the 15 most frequent combinations are shown. **(E)** Normalized Capture-C counts at three selected developmentally regulated genes (promoter baits) and one selected regulatory element (enhancer bait). Gene names and genomic region are indicated above, tissue and stage indicated on the left (colors as in (A)). Arrows indicate regions of interest with significantly differential interaction count between tissues/stages.

The regulatory elements (both enhancers and promoters) targeted for capture were hand-selected from (1) curated databases of *in vivo* validated enhancer elements in transgenic embryos (i.e. Vienna Tiles^29^, REDfly^30–33^ and CAD4^34^) and (2) a curated database of *in situ* hybridisation patterns for thousands of genes (BDGP *in situ* database^35–37^). The initial captures were performed with 637 unique probes. Subsequently, the data from 54 probes was discarded due to either redundant probes capturing the same region or due to low capture efficiency, leaving a final set of 583 bait regions. This represents 303 enhancer baits, 276 promoters, and 4 regions overlapping both enhancers and promoters. The targeted elements were selected primarily based on their dynamic activity in either the myogenic or neurogenic lineages (Fig. 1b and Supplementary Fig. 1a, b). We included baits for 17 gene promoters with important functions during embryonic development, in addition to 6 enhancers that we previously assessed by 4C-seq, serving as internal controls^13^. The expression of this subset of regulatory elements is less exclusive to just one tissue (referred to as the non-exclusive set). We also included 25 baits (9 enhancers, 16 promoters) that have no discernible activity at any of the five tested timepoints or tissues (off-off, e.g. *piwi*), in addition to baits for 8 ‘housekeeping genes’ with ubiquitous activity (on-on, e.g. *Tbp*). The genomic location and activity of all baits are provided in Supplementary Table S1.

The baits were separated into two libraries, predominately targeting either enhancers or promoters, with 26 baits in common to determine the reproducibility and capture efficiency between the two bait pools. Capture-C was performed on two replicates per tissue and time-point (5 sample conditions) with 100 million sorted nuclei per replicate to ensure enough complexity to capture interactions for all regulatory elements. Each sample was independently captured with either the enhancer or promoter bait pools resulting in a total of 20 Capture-C datasets (4 whole embryo 2-3h, 4 Myoblast (Myo) 6-8h, 4 Muscle 10-12h, 4 Neuro 6-8h, 4 Neurons 10-12h). During the FAN sorting, we also actively sorted for non-meso and non-neuro (NM/NN) nuclei representing a mixture of ectoderm and endodermal tissues at both 6-8h and 10-12h, and include this set of 12 Capture-C datasets (4 NM/NN 6-8h, 8 NM/NN 10-12h) as a resource in supplementary material.

Significant interactions above a background distance decay were identified using CHiCAGO^38^, applying a stringent score of ≥ 5, and excluding interacting regions very proximal (<2kb) or distal (>10Mb) to the baits. This identified 52,980 significant interactions across all baits in one or more of the five conditions (2-3h whole embryo (WE), 6-8h Myo, 10-12h Muscle, 6-8h Neuro, 10-12h Neurons), which represents 35,693 unique interactions (Fig. 1c, d; Supplementary Table S2). Analysis of the 26 baits common to both capture libraries revealed high reproducibility in interaction frequencies between replicates both within and between the libraries (Supplementary Fig. 1c-e).

The total number of enhancer and promoter interactions is quite similar at 2-3h (whole embryo) and 6-8h (either Myoblasts or Neuro), ranging from 6000-9500 (Fig. 1c). However, the number of interactions doubles between 6-8h and 10-12h, moving from the stages of cell fate specification to tissue differentiation (Fig. 1c). In the myogenic mesoderm, this changed from 7,157 to 16,388 interactions from 6-8h to 10-12h (Fig. 1c); of these, 11,963 interactions are exclusive to the myogenic lineage (Fig. 1d), 78% (9,281) of which are unique to the later time point of terminal muscle differentiation (Fig. 1d, Muscle 10-12h). Similarly, in the neuronal lineage, there were 6,018 interactions at 6-8h which doubled to 13,907 at 10-12h (Fig. 1c); 9,011 of these are exclusive to the nervous system, with 76% (6,856 interactions) being unique to the later differentiation time-point (Fig. 1d, Neurons 10-12h). The emergence of more enhancer and promoter interactions at 10-12h is also reflected in an increase in the number of interactions per regulatory element (i.e. per bait) at this differentiation stage (Fig. 1d). At 6-8h there was a median of 4 interactions per bait, which increased to over 10 at 10-12h, in both muscle and neuronal tissues (Fig. 1d, with some having more than 100 significantly interacting fragments). There is also an increase in the distance of significant interaction between the 6-8h and 10-12h time points, with more distal interactions appearing in the alter time-point (Supplementary Fig. 1f,g).

The increase in tissue-specific enhancer and promoter interactions during tissue differentiation is nicely exemplified by the loci shown in Fig. 1e. *synapsin* is expressed at late stages in differentiated neurons. Capturing the *synapsin* promoter identifies several significantly interacting regions (loops) that are only present in neuronal cells and only in the later timepoint during terminal differentiation (Fig. 1e, left, arrows). Interestingly, the promoter also loops to a different region at late stages in mesoderm/muscle, a tissue where *synapsin* is not expressed. The gene *Delta* is typically involved in facilitating asymmetric cell fate decisions and has a complex and very dynamic expression in both the developing myogenic and neuronal systems. This is reflected in the observed tissue- and stage-specific *Delta* promoter interactions (Fig. 1e, middle, arrows), suggesting that while *Delta* is active in all tested conditions it employs tissue- and time-specific regulatory topologies. The transcription factor *Zn finger homeodomain 1* (*zfh1*) is also expressed in both tissues at both time-points, yet we only observe very significant changes in promoter interactions at the later stage of terminal differentiation, in both the Muscle and Neurons (Fig. 1e, right, arrows).

In summary, tissue- and stage-specific promoter and enhancer Capture-C revealed hundreds of tissue- and stage-specific looping interactions. While enhancer and promoter interactions displayed only small changes between the early blastoderm and the cell fate specification stage, or between specified myoblasts and the neurogenic system, we observed a large increase in tissue-specific enhancer or promoter loops at the transition between cell-fate specification to terminal tissue differentiation. This suggests that gene regulation during differentiation requires much more *de novo* tissue-specific E-P interactions compared to earlier embryonic timepoints.

### Enhancer-Promoter loops are similar during cell fate specification, but become more specific during terminal tissue differentiation

The examples above indicate that changes in a gene’s expression or in the activity of an enhancer can correlate with changes in their interaction landscape. However, we also observed interaction changes even when the activity of the gene or enhancer does not change, and conversely constant interactions while the activity of the gene or enhancer does change across time or tissues. To investigate this relationship more systematically, we took advantage of the fact that each bait (enhancer or promoter) was hand-selected based on its characterised activity *in vivo* (in both tissue and time), and used this information to categorize all enhancers/promoters (baits) according to their activity dynamics from 2-3h to 6-8h or 6-8h to 10-12h of embryogenesis in both the myoblast/muscle or the developing nervous system (on-off, off-on, on-on, off-off). Correlating these changes in bait (either enhancer or promoter) activity to their changes in topological interactions, revealed that although the overall interaction dynamics within each condition appear very similar, there is a small, but highly significant correlated change in interaction frequencies (Fig. 2a). Enhancers or promoters (baits) that go from off-to-on or on-to-off in their activity have a concordant shift in interaction frequencies going either up or down compared to enhancer/promoters that that do not change (Fig. 2a). There is, therefore, a global trend for enhancer/promoter interactions (as measured by interaction frequency) to mirror the change in activity state of enhancers or promoters. This may reflect new interactions and/or a strengthening or weakening of existing interactions.

**Figure 2:**
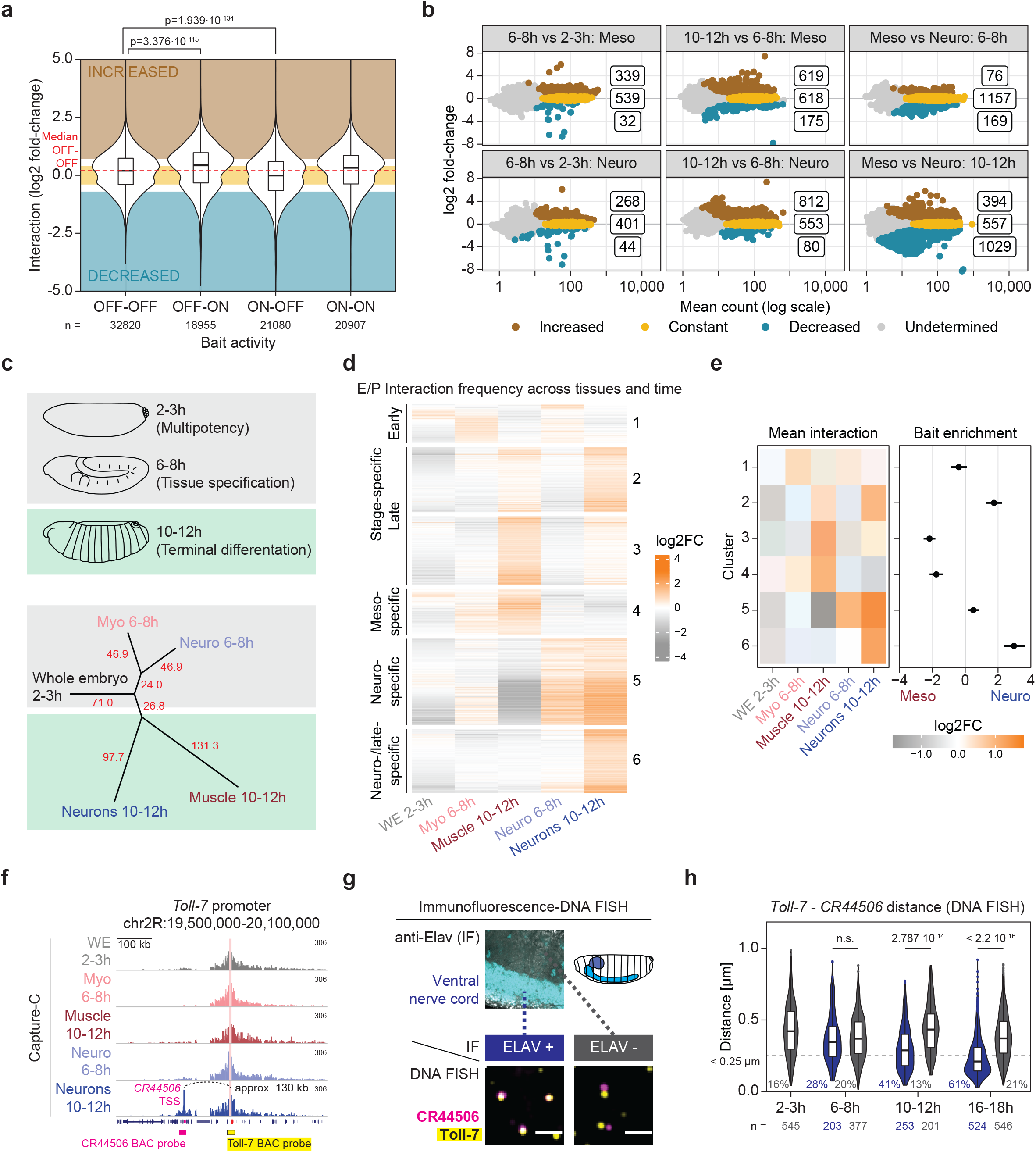
New tissue-specific enhancer or promoter loops are gained during tissue differentiation on top of a largely invariant preformed landscape. **(A)** Violin plot showing changes in interaction frequencies (log2-fold) of significant interacting regions for enhancers/promoters (E/P baits) changing in their activity between conditions (off-off, off-on, on-off, on-on). Number (n) of interacting fragments (CHiCAGO score >5) indicated underneath. P value (above) from non-parametric Wilcoxon test shows a significant concordant trend for increased or decreased interaction frequencies for baits going from off-on or on-off. (B) Scatter plot showing log2-fold change of interacting frequencies between conditions, with significantly increased (brown), decreased (blue) or invariant (yellow) interactions between the two conditions highlighted (numbers indicated). 2782 interactions are significantly differential in one or more of the 6 comparisons, with the majority being found in comparisons involving 10-12h tissues. **(C)** Dendrogram displaying the distance (indicated by branch length) between the 5 developmental conditions based on their enhancer/promoter interaction frequencies for the 2782 differential interactions (from (B)). Myoblasts and neuronal samples are closer to each other at 6-8h and to the 2-3h whole embryo than to their 10-12h tissue counterparts. The two tissues are more divergent in their E/P interactions at 10-12h (indicated by the long branch length). **(D)** K-means clustering (k = 6) of the 2782 differential E/P interactions, that increase (orange) or decrease (grey) relative to the sample average. Most interactions are higher at 10-12h - Clusters 5, 6 in the nervous system and 3, 4 in muscle. **(E)** Enrichment analysis of tissue-specific activity (Neuro vs Meso) of baits (E/P) for the differential interactions within each cluster (shown in (D). Differential interactions increased in a given tissue are generally associated with E/P baits active in that tissue – e.g. clusters with increased interactions in the nervous system (clusters 5, 6) are enriched in E/P baits active in the nervous system. **(F)** Normalized Capture-C counts from a bait, the promoter of *Toll-7* (indicated by red line), of a gene expressed in the nervous system. New interactions are observed at 10-12h in neurons. **(G, H)** Above, immuno stain of Elav expression in the ventral nerve cord. Below, DNA FISH (yellow = *Toll-7*, magenta = *CR44506*) with immuno-stain for a neuronal marker (Elav (cyan)) to assess 3D distance between the indicated genomic loci in the nervous system (Elav+) and adjacent non-neuronal tissue (Elav-) within the same embryos. (H) Box plots of the DNA FISH distance measurements between *Tol-7* and *CR44506* (probe position indicated in (F)) in the nervous system (blue = Elav positive) and non-neuronal tissue (grey = Elav negative). The two loci are significantly closer to each other in neuronal cells compared to non-neuronal cells at 10-12h and 16-18h (while there is not difference at 6-8h). Dashed line =250nm. Number (n) of nuclei measured indicated underneath. P-values from Kolmogorov-Smirnov test.

To examine this further, we identified a stringent set of statistically significant dynamic enhancer/promoter loops (Methods) – retaining differential interactions (FDR 5% and >0.7 log2 fold-change) that were also defined as significant by CHiCAGO in one or more condition (i.e. the regions used in Fig. 2a). This ensures that the distance decay from the bait (implemented by CHiCAGO) is taken into account, while at the same time ensuring that there is a statistically significant quantitative change in interaction frequency between the two conditions (DESeq2). This identified a high-confident set of 4,348 dynamic interactions (2,858 unique) that significantly changed between one or more condition (Fig. 2b, highlighted in brown and blue; Methods, Supplementary Table S2). We also tested for the opposite – interacting regions that do not change significantly (FDR 5% and <0.4 log2 fold change, Methods) – defining a stringent set of 6,853 ‘invariant’ interactions (3,164 unique) in one or more condition (Fig. 2b, highlighted in yellow, Supplementary Table S2). Again, we confirmed the reproducibility of the data by comparing the log2 fold-change between the two capture libraries using the 26 baits common to both (Supplementary Fig. 1e).

In line with our observations above, there are very few differential interactions between 2-3h, the multipotent blastoderm stage, and 6-8h during the initial specification of myoblasts and neuroblasts: 371 and 312 interactions comparing 2-3h to Myo or Neuro 6-8h, respectively (Fig. 2b). The majority of these are gains in interactions at 6-8h; 91% Myo (339/371) and 86% Neuro (268/312). Conversely, the number of significantly constant (tissue invariant) interactions between Myo and Neuro cells at 6-8h is far greater than the number of differential loops between the two cell types (Fig. 2b). These findings are similar to what we observed previously assessing 100 enhancers at the same two stages of embryogenesis by 4C-seq, where less than 8% of interacting regions changed^13^. This indicates that despite the large transcriptional changes occurring during these stages of embryogenesis, which includes zygotic genome activation, the segmentation and gastrulation of the embryo, and subsequent cell fate specification events within the myogenic and neuronal lineages, many embryonic enhancers are already in 3D topologies at the early blastoderm stage, which appear to be maintained throughout embryogenesis.

The picture becomes very different comparing the stages of cell fate specification (6-8h) to the later developmental stages of terminal tissue differentiation (10-12h). Comparing regulatory interactions within a tissue between these two time-points revealed a substantial increase in the number of differential interactions: 794 in the myogenic and 892 in the neuronal lineages between 6-8h and 10-12h (Fig. 2b, brown and blue). The majority of these are again increased interactions at the later timepoint (78% [635/794] Muscle, 91% [844/892] Neurons), indicating that many enhancer-promoter loops are formed or strengthened during tissue differentiation and that they are added on top of a pre-existing regulatory topology (Fig. 2a, brown). As a consequence, the number of shared (invariant) interactions between the two tissues is lower at 10-12h (during terminal differentiation) compared to 6-8h (during specification of Myoblasts and the neurogenic system: 557 vs 1157 (Fig. 2b, yellow). Both results indicate more diversification in enhancer/promoter interactions between tissues during differentiation (at 10-12h), while there is more similarity at earlier stages. We confirmed this in an independent analysis, taking the normalised interaction counts from all differential interactions in one or more comparison (Methods), and constructing a dendrogram to observe the relative distance between the interactions counts at each condition. This indicates that interactions with enhancers and promoters are more similar between Myoblasts and the neurogenic system at 6-8h (when the cells are specified), than they are within a tissue between the two time-points, i.e. between Myoblasts (6-8h) and Muscle (10-12h) or Neuro (6-8h) and Neurons (10-12h) (Fig. 2c). The longer branch length at 10-12h, indicates that these tissue-specific enhancer or promoter loops become more divergent at later stages of a tissue’s development. The majority of differential interactions are present in only one or two comparisons indicating their high tissue- and/or stage-specificity (Supplementary Fig. 2a).

Although we note, we may be underpowered to detect loss of interactions, as more of the baits (enhancers and gene promoters) become activated, or remain active, rather than inactivated at the later developmental timepoint (Fig. 1b).

Changes in the strength (interaction frequency) of these differential interactions is very apparent after unsupervised (K-means) clustering, which revealed three broad classes (Fig. 2d). A fraction of dynamic interactions is time-rather than tissue-specific, being present during tissue specification (6-8h) or at terminal differentiation (10-12h) (Fig. 2d clusters 1-3). However, the majority of differential loops are tissue-specific (either in the muscle or neuronal system), and get stronger in, or are specific to, the later timepoint during tissue differentiation (10-12h) (Fig. 2d, Muscle (cluster 4) and Neurons (clusters 5, 6)). The bulk of the interactions within each of these clusters are also only significant (CHCAGO score 5) in the appropriate tissue-differentiation condition (Supplementary Fig. 2b). This provides strong evidence that these differential enhancer or promoter interactions are truly *de novo* interactions in that tissue or developmental time-point.

To assess the relationship between these differential interactions and E/P activity, we next determined if they preferentially occur in the tissues in which the corresponding baits are active. This is indeed the case; enhancers or promoters (baits) with activity in the myogenic mesoderm or muscle are significantly enriched in clusters with strongest interactions frequencies (differential contacts) in Myoblast or Muscle cells (Fig. 2e, clusters 3, 4), while baits with activity in the nervous system are enriched in interactions that are strongest in the nervous system (Fig. 2e, clusters 2, 5, 6; Supplementary Fig. 2c). This significant enrichment indicates that tissue-specific E/P loops generally occur in the cell type where the enhancer or promoter is active, again linking regulatory element interaction to activity.

For example, the promoter of the *Toll-7*, a gene expressed in the ventral nerve cord, has a specific loop in neuronal (but not muscle) cells during terminal differentiation at 10-12h and not at the early developmental stage (Fig. 2f, g). This neuronal-specific loop involves the *Toll-7* promoter and the promoter of a long non-coding RNA gene (lncRNA *CR44506*) approximately 130 kb away (Fig. 2f). We verified the proximity of these loci by DNA FISH combined with immunofluorescence to visualise the nervous system (Elav) in developing embryos (Fig. 2g). This confirmed that the *Toll-7* promoter is in closer proximity to the interacting lncRNA gene’s promoter specifically in the nervous system (Elav+ cells), compared to adjacent non-neuronal cells (Elav-) (Fig. 2h). While there is no significant difference between the distance of these loci in neuronal versus non-neuronal cells at 6-8h, both regions come into proximity specifically at 10-12h in neurons confirming the Capture-C results, with an even higher fraction of cells showing close proximity at the end of embryogenesis (16-18h): 61% of nuclei have these loci within 250nm or closer (Fig. 2h).

This exploration of the relationship between enhancer and promoter activity (i.e. when and where both are active) during embryogenesis and their 3D proximity (i.e. specific looping) revealed three general properties. First most enhancers and promoters that are active at 6-8h, spanning stages when myoblasts and the neurogenic system are being specified, are already in predefined topologies earlier in embryogenesis, at the blastoderm stage, as we observed previously for a subset of enhancers. Second, more tissue-specific interactions occur at later embryonic stages, during terminal tissue differentiation. Third, we generally observe gains in new interactions (or strengthening) rather than losses (or weakening), in specific tissues during differentiation. This indicates that regulatory topologies build on a foundation that is established during early stages of embryogenesis, which is then added to by additional tissue-specific loops during tissue differentiation.

### Binding of well-characterised insulators can only explain a small fraction of differential interactions between enhancers and promoters

Insulator proteins can facilitate chromatin loops that either enable enhancer-promoter interactions or block them, depending on the relative location of the insulator’s binding site with respect to the enhancer and promoter. In addition to CTCF, *Drosophila* has several other zinc finger proteins that bind to insulator elements either directly or indirectly. To determine which insulator proteins (or combinations of them) might be involved in differential or invariant enhancer and promoter interactions during embryogenesis we searched for co-occupancy of different insulators. To facilitate this, we first generated tissue- and stage-specific ChIP datasets matching our Capture-C data. Mesodermal and neuronal nuclei were FANS purified from staged embryos at 6-8h and 10-12h, as used for the Capture-C (Fig. 1a), and used for ChIP-seq on Su(Hw), BEAF-32 and CTCF, representing 4 conditions, each with biological replicates.

As this is the first tissue-specific view of insulator binding during *Drosophila* embryogenesis, we initially explored the dynamics and tissue-specific differences between insulator protein occupancy in myoblast/muscle and neuro/neurons. Each factor binds to thousands of regions, with 5,359, 5,071 and 3,819 significant peaks (FDR 0.05) identified for Su(Hw), BEAF-32 and CTCF, respectively, at one or more condition(s) (Fig. 3a, Methods). 2,838 regions are bound by two or more insulator proteins (within 50bp) and 1,359 by all three factors, although not necessarily at the same timepoint or tissue (Fig. 3A, Supplementary Tables S4-6, Methods). The majority of regions bound by BEAF-32 and Su(Hw) are not bound by other factors (within a 50 bp window), while CTCF has the most combinatorial binding; ∼70% [2,685/3,819] of all CTCF peaks are bound by more than one factor (Fig. 3a). An example is shown at the *even skipped* locus where the gene’s rather compact regulatory landscape (as observed by Capture-C profile) is flanked by insulator binding in both tissues and time-points (Fig. 3b); CTCF and Su(Hw) on the left and BEAF-32, CTCF and Su(Hw) on the right. Interestingly, the occupancy at six functionally characterized insulator elements is more variable across tissues and time-points, especially in the nervous system at 6-8h and 10-12h (Supplementary Fig. 3a).

**Figure 3:**
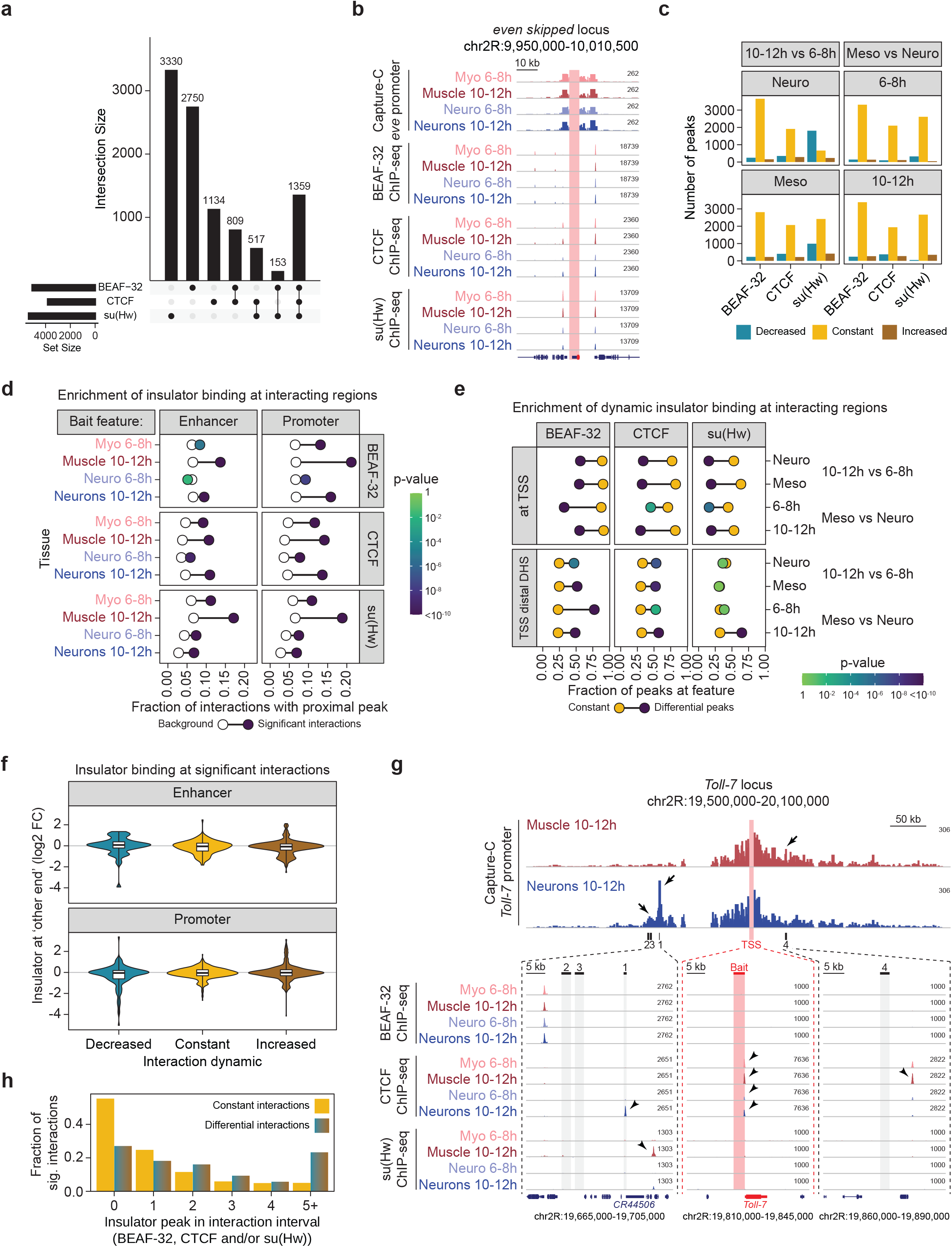
Differential insulator binding accounts for a small fraction of differential bait interactions. **(A)** UpSet plot of significant insulator ChIP-seq peaks in the indicated four conditions. **(B)** Normalized Capture-C and insulator ChIP-seq signal in the indicated conditions at the *even skipped* locus, bait (*eve* promoter) highlighted by red bar. **(C)** Bar chart of the number of constant (yellow), decreased (turquoise) and increased (brown) insulator peaks in the respective comparisons. With the exception of Su(Hw) in the neuronal tissue, insulator peaks are largely invariant between tissues and across timepoints. **(D)** Enrichment analysis of all insulator peaks in proximity to capture C interacting regions. The frequency of insulator peaks in proximity (<500bp) to ‘otherEnd’ significant (colored dots) and non-significant (white dots) interactions in the respective samples. The shade color of colored dot denotes the p-value (Fisher exact test). Data for the promoter and enhancer baits are shown separately. CTCF and Su(Hw) are enriched at the ‘otherEnd’ of both promoter and enhancer baits, while BEAF-32 is more enriched at interacting regions of promoter baits. **(E)** Enrichment of differential insulator peaks - frequency of constant (yellow dot) and differential (blue/green dot) insulator peaks (between time/tissues) in proximity (<500bp) to the ‘otherEnd’ of significant capture C interactions, separated by promoter and enhancer baits. Shade of the blue/green dot indicates the significance (Fisher exact test) of frequency differences between constant and differential insulator peaks. For all 3 insulator proteins, constant/invariant peaks are significantly more enriched at promoters, while differential BEAF-32 and CTCF peaks are significantly more frequent at distal DHS. **(F)** Violin plots showing quantitative changes in insulator binding (log2 fold change (FC) ChIP-seq signal) at ‘otherEnds’ (<500bp) of Capture C baits with decreased (blue), constant (yellow) or increased (brown) interactions across the same conditions (tissue or time). Changes in enhancer/promoter interactions is generally not correlated with changes in insulator binding. **(G)** Normalized Capture-C counts at the *Toll-7* locus (promoter bait) in Muscle and Neurons at 10-12h. Below, zoom-in showing insulator ChIP binding in the indicated windows. Differential neuronal-specific interaction between *Toll-7* promoter and *CR44506* at 10-12h is associated with differential binding of CTCF in Neurons at 10-12h. Lack of interaction in Muscle 10-12h is associated with Su(Hw) binding in Muscle between *Toll-7* and *CR44506*. A muscle specific 10-12h interaction between the *Toll-7* promoter and *Toll-7* #4 enhancer coincides with muscle-specific CTCF adjacent binding. **(H)** Number of differential insulator peaks (x-axis) in the interaction interval (between bait and otherEnd’ +/-5kb) for both constant and differential capture C interactions. Differential insulator peaks are found more frequently in the interaction interval of differential interactions than constant interactions.

Examining this more globally, almost half (44% (4429/10052)) of insulator peaks (for any of the 3 factors) have a statistically significant change (FDR 5% and >0.7 log2 fold-change) for any one of the factors between tissues (e.g. Myo vs Neuro at 6-8h) or between time-points within a tissue (e.g. Myo 6-8h vs Muscle 10-12h) (Supplementary Fig. 3b, Methods). This indicates a roughly equal split between dynamic (44%) and more invariant (56%) binding of these factors between time-points or tissues, at these stages of embryogenesis (Fig. 3c). The most changes are observed for Su(Hw), where 1959 peaks have reduced binding at the later timepoint (10-12h) in either the neuronal (1809 peaks), mesodermal (991) or both (841 peaks) tissues (Fig. 3c, Supplementary Fig. 3b). Differential insulator peaks also show less combinatorial binding with the other two insulator proteins (Supplementary Fig. 3c).

To assess whether these insulator proteins could facilitate enhancer and promoter interactions, we first determined if there is a general enrichment in the binding of the three insulator proteins within 500bp of the ‘other end’ of the significant enhancer or promoter loops (CHiCAGO peaks, Methods). This revealed a general higher likelihood for insulator binding at interacting regions, with the most significant enrichments at later stages during differentiation (Muscle, Neurons 10-12h). Beaf-32 and Su(Hw) are particularly enriched in Muscle 10-12h, being about 4 times higher at promoter interacting regions compared to background (Fig. 3d). Across the entire dataset, 21% (3879/18608) of all enhancer and 26% of all promoter (6269/24448) interacting regions are bound by one or more of the three insulators in the same tissue.

Comparing the location of constant (invariant) versus dynamic insulator binding revealed an interesting split (Fig. 3e): Constant peaks are highly enriched at interacting promoters (<500bp of a TSS), a trend that holds true for all three insulators in all comparisons, while dynamic insulator binding is enriched at distal interacting DHS (putative enhancers), which is particularly strong for BEAF-32 in Neuro vs Myo at 6-8h (Fig. 3e). However, comparing the quantitative changes in insulator binding to changes in interaction frequencies revealed surprisingly little general concordance between the changes in enhancer or promoter interactions across time or tissue (Fig. 3f).

However, there is a small subset of regions with correlated changes in insulator binding and enhancer-promoter loops, including the interactions we confirmed by DNA FISH at the *Toll-7* locus. The neuronal-specific proximity of the lncRNA gene *CR44506* promoter and the *Toll-7* promoter at 10-12h, for example, is accompanied by a CTCF peak, specifically at 10-12h in neurons (Fig. 3g, Toll-7 #1). Similarly, there is mesoderm-specific CTCF binding adjacent to the Toll-7 #4 element, specifically interacting in the mesoderm (Fig. 3g, #4). This context-specific insulator binding is similar to previous observations at the *Ubx* locus, where CTCF binds close to an enhancer specifically in tissues where *Ubx* is expressed (3^rd^ leg imaginal disc) and not in tissues where it is repressed (1^st^ leg imaginal disc)^39^. In that context, the CTCF site was close to a Su(Hw) bound gypsy element, which interfered with CTCF binding leading to a *Ubx* mis-expression phenotype, suggesting that 3^rd^ leg imaginal disc specific binding of CTCF facilitates an instructive tissue-specific loop between the *Ubx* enhancer and promoter. Interestingly, we also observe a Su(Hw) bound region ∼10 kb downstream of Toll-7 #1 specifically in Muscle 10-12h (Fig. 3g, left panel), suggesting that it perhaps acts in a similar manner to block a *CR44506*-*Toll-7* interaction in muscle cells.

Similarly, the neuronal gene *chinmo* (*chronologically inappropriate morphogenesis*), interacts with a putative neuronal enhancer (chinmo #1) specifically in neurons at late stages (Supplementary Fig. 3d). This region has decreased Su(Hw) binding in the late nervous system and mesoderm, while Su(Hw) occupancy in this region is maintained in Muscle at 10-12h, suggesting that this again may be blocking loop formation in these cells. At the *robo3* (*roundabout 3*) locus, there is a CTCF peak close to the *robo3* promoter and an interacting putative enhancer (*robo3* #1) in Neurons (but not the Muscle) cells at 10-12h, and not at 6-8h (Supplementary Fig. 3e), suggesting that insulator binding at one end of the loop might be involved in its regulation. There is a concomitant decrease in intragenic Su(Hw) binding downstream of the *robo3* promoter, in addition to changes in BEAF-32 and CTCF binding close to the promoter in Neuro cells at 10-12h when this looping interaction is formed (Supplementary Fig. 3e).

In addition to functioning by binding close to enhancers and promoters, insulators can also function when placed in between, at least in transgenic reporter assays. We therefore also examined differential insulator binding in the intervening sequence between a differential enhancer-promoter interaction, similar to the differential Su(Hw) peak several kilobases downstream from the *CR44506* interaction with the Toll-7 promoter (Fig. 3g, #1). Each interaction was scored based on the number of differential insulator peaks in the intervening sequence between two interacting fragments as well as 5 kb up- or down-stream of the bait and other-end (referred to as interaction interval). A substantial fraction of differential interactions (71%), have at least one differential insulator binding peak within their interaction interval, which is higher than that observed for invariant interactions (44%, Fig. 3h). Moreover, the density of differential insulator peaks per kb was significantly higher within the interaction interval for differential enhancer or promoter interactions compared to invariant interactions (p-value 6.6×10^−88^ Mann-Whitney test, Supplementary Fig. 3f).

However, there was no global concordance between the direction of change of the insulator peak and changes in interaction frequency.

Taken together, although insulator binding is generally enriched at the anchors of significantly interacting regions, changes in insulator binding is not globally correlated with changes in enhancer or promoter interaction frequency, at least for the insulators examined here. Therefore, although differential insulator binding may play a role at some loci and have an impact on differential E-P interactions, as we show for CTCF, such context-specific binding (at least for these three insulators) can only explain a fraction of the differential loops we observe during tissue differentiation (compared to early developmental stages) or between the mesoderm and nervous system.

### Dynamic enhancer and promoter loops are correlated with dynamic features of regulatory activity

To determine how these differentiation loops might be regulated, we examined if they are associated with changes in chromatin accessibility or chromatin modifications. Each significant interaction, whether it is differential or constant, is defined by two ‘anchor points’: the bait (either an enhancer or promoter) and the ‘other end’, which can include enhancers, promoters and regions of unknown function (Fig. 4a, c). Interaction frequency presumably depends on molecular events occurring at one or both of these ends, and potentially also loci between and adjacent to the two anchor points.

**Figure 4:**
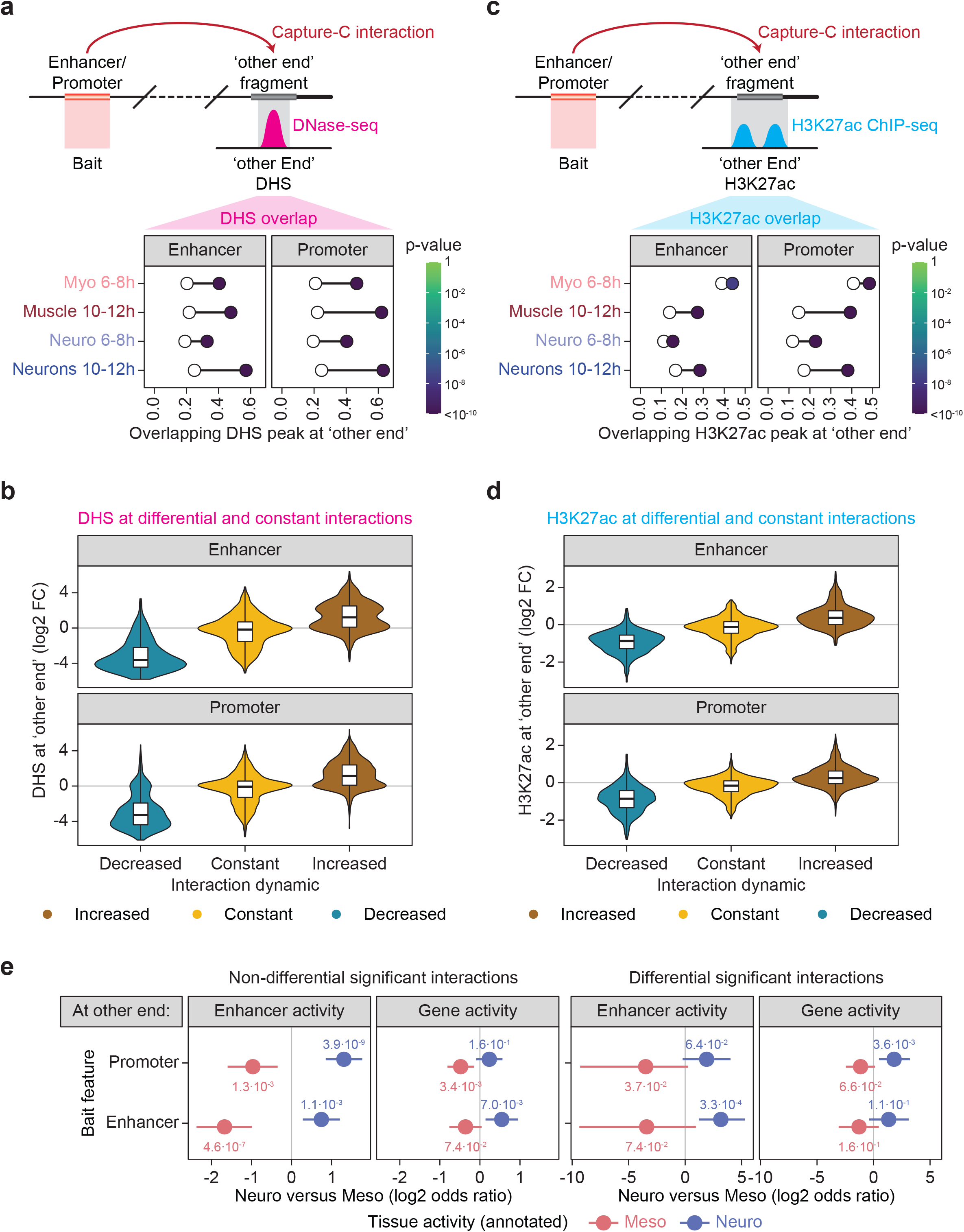
Changes in enhancer/promoter interactions have concordant changes in chromatin accessibility and chromatin state. **(A)** Enrichment of DHS in proximity to the ‘otherEnd’ of all significant interactions (shown in the schematic above). Plot shows frequency of DHS within 500bp of the ‘otherEnd’ in the respective condition, for all significant (colored dot) or non-significant (white dot) interactions. Shade of colored (dot) indicates p-value (Fisher exact test). Enrichments for enhancer and promoter baits shown separately. DHS are more frequently at the ‘otherEnd’ of significant interactions at 10-12h samples, compared to 6-8h. **(B)** Violin plots showing quantitative changes in DHS signal (log2 fold change (FC)) at ‘otherEnds’ (<500bp) of Capture C baits with decreased (blue), constant (yellow) or increased (brown) interactions across the same conditions (tissue or time). The direction of change in DHS signal correlates with change in interaction frequency. **(C)** Similar to (A), enrichment of H3K27ac peaks within 500bp of the ‘otherEnd’ of all significant interactions. Plot shows frequency of H3K27ac within 500bp of the ‘otherEnd’ in the respective condition, for all significant (colored dot) or non-significant (white dot) interactions. Shade of colored dot indicates p-value (Fisher exact test). **(D)** Similar to (B), violin plots showing quantitative change in H3K27ac signal (log2 FC) at ‘otherEnds’ (<500bp) of Capture C baits with decreased (blue), constant (yellow) or increased (brown) interactions across the same conditions (tissue or time). The direction of H3K27ac change correlates with changes in interaction frequency, especially at 10-12h. (E) Enhancer/Promoter baits interact with enhancers or promoters that are active in the same tissue (either neuro or myo/muscle). Baits active in both tissues were excluded. Enrichment (log2 odds ratio) of features active in the two tissues indicated on the x-axis. A positive value indicates enrichment in baits with Neuro activity (relative to Muscle), negative values indicate enrichment in Muscle activity (relative to Neuro). Dot indicates the observed log2 odds ratio, whiskers show the 95% confidence interval. *Left*, enrichments for non-differential enhancer/promoter interactions. *Right*, enrichments for differential E/P interactions. Enhancer/promoter baits (both differential and constant) active in one tissue preferentially interact with genomic features active in the same tissue.

We first assessed the relationship between changes in open chromatin and changes in interaction frequency, assessing concordance or discordance of differential interactions going from on-to-off or off-to-on and invariant interactions. We focused on the timepoints of tissue specification (6-8h) and terminal differentiation (10-12h), as more differential interactions are occurring at the later time-point. For this we used our previously generated tissue- and time-specific DNase hypersensitivity site (DHS) data at matching tissues and developmental timepoints^28^. Interacting regions at the ‘other end’ are generally highly enriched in DHS peaks, suggesting interactions between our enhancer and promoter baits with other regulatory elements (Fig. 4a; Supplementary Table S3). The overlapping DHS fraction increased at the later developmental timepoint in both tissues, and was similar for both the enhancer and promoter bait sets (Fig. 4a). As DHS peaks are highly enriched in transcription factor binding, this indicates that developmental enhancers and promoters preferentially interact with regions bound by, or at least accessible to, transcription factors.

Even more striking, there is a high concordance between the changes in enhancer/promoter looping interactions and the changes in DHS peaks at the ‘other end’, where increasing interacting frequency in a given tissue is correlated with increasing DHS signal at the ‘other end’ in that tissue and vice versa (Fig. 4b, brown, blue). Conversely, invariant interactions are not associated with changes in DHS between conditions (Fig. 4b, yellow). This is consistent with a gain or loss of TF binding in the myogenic or neuronal linages underlying increased or decreased enhancer and promoter interactions in those tissues. This is in sharp contrast to what we observed with changes in insulator binding, which are generally not correlated with changes in E-P interaction frequency (compare Fig. 3f to 4b).

To identify potential regulators of tissue-specific (or developmental stage-specific) enhancer-promoter loops, we performed motif analysis at the ‘other End’ of enhancer and promoter interactions, using the CIS-BP database of *D. melanogaster* binding motifs^40^.

Using our previously published DHS data from the same tissues and stages of embryogenesis^28^, we searched for enriched motifs within DHS regions proximal (<500bp) to the ‘other end’, for all interacting regions and separately for differential interacting regions (Supplementary Fig. 4, Methods). This identified 20 significantly enriched motifs for all interacting regions (adjusted p-value < 1×10^−4^), including motifs for factors known to play a role in controlling enhancer-promoter communication or chromatin topology e.g. Trl/GAF^41– 43^ and Clamp^44,45^, as well as a number of factors that have not been directly implicated in genome topology before (Supplementary Fig. 4a). Motifs enriched specifically at the ‘other end’ of differential loops (compared to DHS of non-differential interactions) identified motifs for 7 factors (Supplementary Fig. 4b). This includes motifs for several transcription factors essential for the development of the respective tissues, including Mef2, enriched at the ‘other end’ of Muscle (compared to Neuron) specific interactions at 10-12h, and Ttk and l(3)neo38 enriched at Neuronal (compared to Muscle) specific interactions at 10-12h. These enrichments are against a background of tissue-matched DHS and suggest that lineage-specific transcription factors may also play a role in regulating loop formation, either directly or via the activation of developmental enhancers, which then form a loop.

As differential loops generally occur in the tissues and time-points where the enhancer and promoter baits are active (Fig. 2e), it suggests that many differential interactions may be regulatory in nature. To assess this, we used H3K27ac as an indicator of active enhancers and promoters and looked for changes in activity within a tissue across the two time-points.

To facilitate this, we performed tissue- and stage-specific ChIP-seq for H3K27ac with sorted nuclei matching the tissues (Muscle and Neuronal) and stages (6-8h and 10-12h) analysed by Capture-C (Fig. 1a). Similar to DHS, H3K27ac is generally enriched at the interacting regions’ ‘other end’, particularly at the timepoint of terminal differentiation (10-12h), suggesting that the enhancer and promoter baits are interacting with regions that are either active promoters or enhancers (Fig. 4c; Supplementary Table S7). These enrichments for both DHS and H3K27ac at the loop anchors are significantly higher at the later time-point compared to background (Fig. 4a, c, compare filled to white circles). Importantly, differential E-P interactions between any two conditions are associated with concordant changes in H3K27ac signal (Fig. 4d): a gain or loss of interactions from enhancers or promoters are associated with a gain or loss in the activity status of the genomic element at the ‘other end’ (Fig. 4d, brown, blue). Conversely, invariant interactions have no change in H3K27ac signal (Fig. 4d, yellow). This again indicates that changes in E/P interactions are associated with changes in activity confirming our previous finding (Fig. 2e), using an independent, more global and quantitative metric of enhancer activity.

A fraction of the interacting ‘other end’ overlap characterised embryonic enhancers or promoters and/or genes. Examining the expression of these genes or the activity of the enhancers shows that they preferentially interact with baits (enhancers or promoters) that are active in the same tissue (Fig. 4e, Methods). For example, enhancers and genes active in myoblasts or muscle are enriched at the ‘other end’ of myogenically active baits (compared to neuronally active baits) (Fig. 4e, right, red). Likewise, neuronally active genes and enhancers are enriched at the ‘other end’ of loops from neuronally active baits (compared to Myo/Muscle active baits) (Fig. 4e, right, blue). This enrichment is higher and more significant when the looped ‘other end’ is an enhancer compared to a gene, irrespective of whether the bait is itself a gene promoter or enhancer. This indicates that dynamic (or tissue-specific) interaction loops from an enhancer or promoter interact with other elements (enhancers and promoters) that are active in the same tissue and time-point, suggesting that they could have regulatory function. Interestingly, we note that the same enrichments also hold true for all remaining non-differential interactions (Fig. 4e, left), suggesting that many invariant interactions (constitutive loops) may also have underlying regulatory function. This extends previous findings that some topologies are pre-formed or more invariant, while others are more condition (tissue- or stage-) specific. Our results suggest that genes active later in embryogenesis use a mixture, where they have pre-formed topologies early in embryogenesis that are added to later by *de novo* interactions during tissue differentiation.

Taken together this suggests that differential loops coming from enhancers or promoters have regulatory properties (i.e. concordance in their changes in chromatin accessibility, chromatin state and tissue activity) suggestive of being functionally important for gene regulation. These interactions are therefore good candidates for functional analyses, as we have done below. However, it is important to bear in mind that many constitutive (invariant) interactions may also have regulatory functions as the loops ‘other ends’ are also enriched in DHS, H3K27ac and matching tissue activity.

### Concordant changes in interaction frequency and chromatin state across conditions is a good indicator to connect functional enhancer-promoter pairs

Given the general concordance between differential tissue-specific loops and tissue-specific changes in both the bait’s (enhancer or promoter) and the interacting loop other end’s activity, described above, we reasoned that a combination of all three (i.e. correlated changes in interactions, accessibility and H3K27ac) might identify functional enhancer or promoter pairs. To assess this, we selected 12 loci (promoter baits) with different properties, and determined if their interacting regions (19 in total) could function as enhancers *in vivo*, and if they recapitulate part of the gene’s expression (Fig. 5, Supplementary Fig. 5 and Table S8).

**Figure 5:**
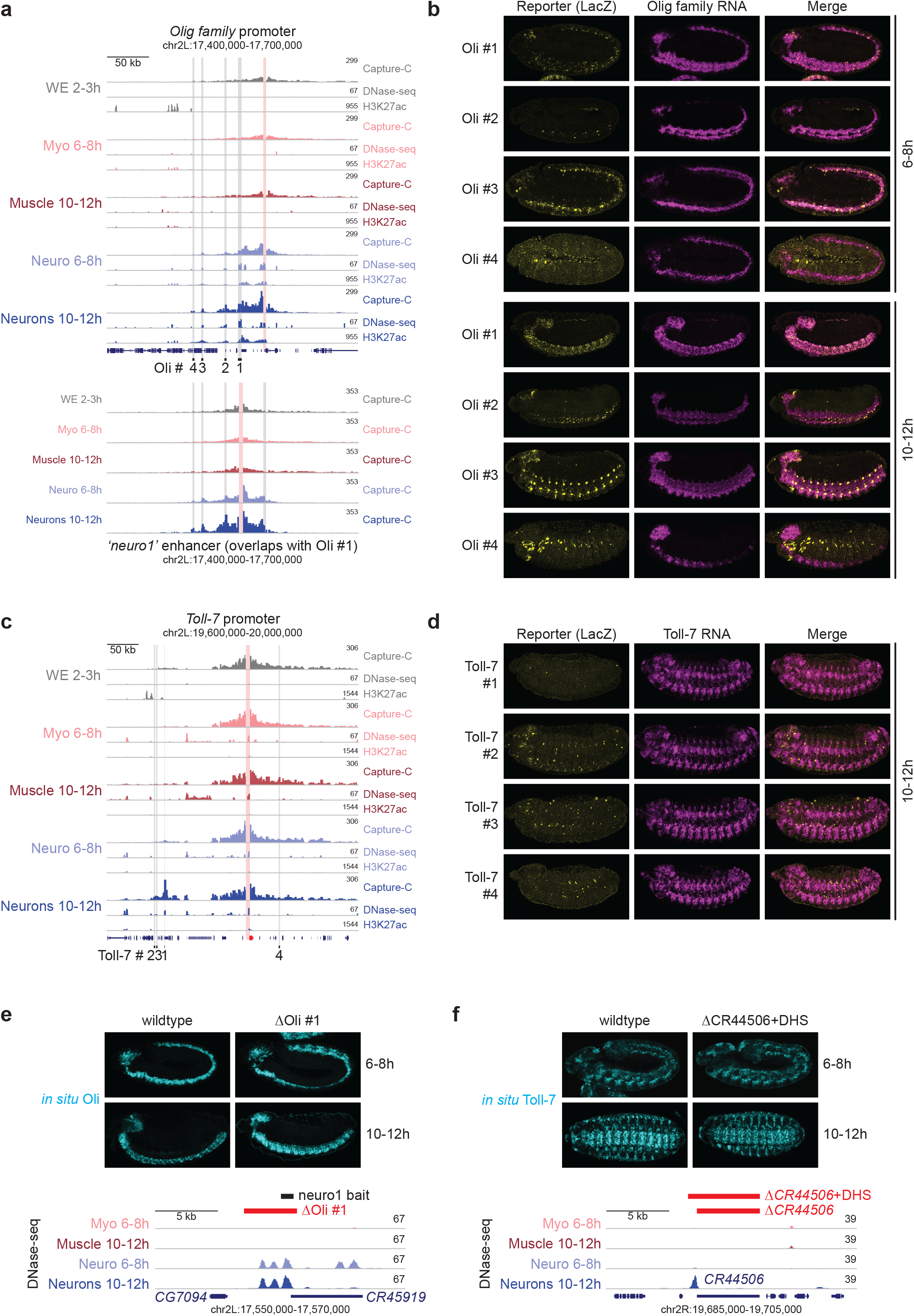
Differential Capture-C interactions combined with changes in chromatin state often represent functional enhancer elements. **(A)** *Upper*: Normalized Capture-C, DNase-seq and H3K27ac ChIP-seq signal at the *Olig family* (*Oli*) locus in 5 conditions. Vertical red bar = bait (*Oli* promoter), grey bars = position of interacting regions tested for enhancer activity. *Lower*: Reciprocal Capture C from the *neuro 1* (oli #1) enhancer (bait = vertical red bar). Normalized Capture-C counts in 5 different conditions. Significant interactions between neuro1 and the *Oli* promoter and other elements indicated by grey bars. Region deleted (neuro1) by CRISPR/Cas9 is indicated (red horizontal bar). **(B)** Double fluorescent *in situ* hybridization of embryos for four transgenes testing enhancer activity of *Oli* #1-4 at the indicated stages. Yellow = reporter RNA (*lacZ*), magenta = *Oli* RNA. Oli #1-3 function as enhancers overlapping part of *Oli* expression at 10-12h. **(C)** Normalized Capture-C, DNase-seq and H3K27ac ChIP-seq counts at the *Toll-7* locus in 5 conditions. Vertical red bar = bait (*Toll-7* promoter), grey bars = position of interacting regions tested for enhancer activity. **(D)** Double fluorescent *in situ* hybridization of embryos for four transgenes testing enhancer activity of *Toll-7* #1-4 at the indicated stages. Yellow = reporter RNA (*lacZ*), magenta = *Toll-7* RNA. Toll-7 #2-4 have sporadic enhancer activity is a small subset of cells. **(E)** RNA *in situ* hybridization of *Olig family* expression in wildtype embryos and *Oli* #1 deletion (∆Oli #1 (neuro1)) homozygous embryos. *Below*, zoom in of the deleted region showing the location of *CR45919* and 3 neuronal DHS peaks. **(F)** RNA *in situ* hybridization of *Toll-7* expression in wildtype embryos and Toll-7 #1 deletion (∆Toll-7 #1) homozygous embryos. *Below*, zoom in of deletions showing the location of *CR44506* and a neuronal 10-12h specific DHS peak.

The *Olig family* (*Oli*) gene is expressed in the brain and central nervous system (CNS) and required for motoneuron axon pathfinding and larval and adult locomotion^46^. A bait from the *Oli* promoter interacts with a number of upstream regions specifically in neuronal cells – Oli #1-3 from 6-8h onwards and a more distal region (Oli #4) at 10-12h (Fig. 5a, upper). Oli #1 (called ‘neuro1’, based on tissue-specific accessibility from^34^) was included in our enhancer bait set and we detect a reciprocal interaction with the *Oli* promoter and the other three putative regulatory elements (Oli #2-4), confirming these interactions (Fig. 5a, lower). All regions (with the exception of the more distal one; Oli #4) have a DHS and H2K27ac peak in neuronal cells, and these peaks (and their looping interactions with the promoter) are not observed in muscle cells (Fig. 5a). These regions are therefore prototypical examples of tissue- and stage-specific looping, being only present in the tissue where the gene is expressed. Of note, Oli #1 & #2 contain Ttk motifs in the underlying DHS, while the *Oli* promoter has increased CTCF binding in neuronal cells and a reduction of Su(Hw) binding adjacent to the Oli #1 element (data not shown). We tested all four regions for enhancer activity in transgenic reporter assays assessing their activity at all stages of embryogenesis (Fig. 5b). Three of the four regions have neuronal enhancer activity overlapping the expression of the *Oli* gene at the appropriate stages of embryogenesis, confirming that these regions are neuronal enhancers and suggesting that their interactions with the promoter are instructive, coinciding with the gene’s expression. The activity of the enhancers is also partially overlapping each other (e.g. in the ventral nerve cord) suggesting that some of these enhancers are likely to be partially redundant. The most distal region (Oli #4) showed no significant overlap with *Oli* expression, which is also the region that had no H3K27ac signal.

In the *Toll-7* locus, we tested four regions interacting with the gene’s promoter, three of which (Toll-7 #1-3) have neuronal specific and one (Toll-7 #4) has muscle specific DHS at 10-12h^28^, matching the expression of the *Toll-7* gene (Fig. 5d). However, none of the four regions have significant H3K27ac peaks. Three regions (Toll-7 #2-4) showed weak enhancer activity that overlaps with *Toll-7* expression in a small subset of cells: Toll-7 #2 in the ventral nerve cord (VNC), #3 in endoderm and a subset of VNC and visceral muscle cells, #4 in a small subset of cells in VNC and visceral muscle (Fig. 5e). The fourth region, Toll-7 #1, which is the most significant differentially interacting region with the Toll-7 promoter, has no enhancer activity (Fig. 5d, e), and overlaps the promoter of the lncRNA *CR44506*. We confirmed that the proximity between the *Toll-7* promoter and *CR44506* is specific to the nervous system and is not present in non-neuronal tissues by DNA FISH, as described above (Fig. 2f-h). *CR44506* is very weakly expressed in the nervous system in a pattern that partially overlaps Toll-7 at the stages of embryogenesis when the interaction is detected (10-12h, Supplementary Fig. 5a), suggesting a possible link between their co-expression and physical proximity. However, very few cells had detectable nascent RNA signal that overlapped, although this could be due to the low expression of the lncRNA. Alternatively, Toll-7 #1 could function as a structural element, facilitating interaction of the *Toll-7* promoter with nearby lociWe al so tested two regions interacting with the *Delta* promoter, a more proximal region (Dl #1) with specific interactions in muscle and a more distal region (Dl #2) with neuronal specific interactions. Both regions have no obvious association with our tested set of insulators, Dl #2 has an underlying Ttk motif. The mesodermal interacting region (Dl #1) had enhancer activity in the visceral and somatic muscle, which overlaps *Delta* expression in these tissues. The distal element (Dl #2) also had enhancer activity but in the endoderm and visceral muscle, two tissues that do not match the enhancers neuronal specific interactions (Supplementary Fig. 5b).

In total, of the 19 interacting elements tested for enhancer activity *in vivo* (Supplementary Table S8); *Oli* (4 test regions), *Toll-7* (4), *Dl* (2), *lmd, bap, tin, robo3, hkb, VAChT, danr, chinmo, Dop1R1*, 14 (74%) showed enhancer activity in the same cell type as their interacting promoter, and at least partially overlap the expression of the interacting gene, e.g. *Dl* #1 in late muscle, *lmd* in a subset of somatic muscle (Supplementary Fig. 5b, c). For some elements, the enhancer activity was rather weak and limited to a small subset of cells in the respective tissue e.g. the *robo3* element was active in only a few expressing cells of the embryonic brain (Supplementary Fig. 5d), while for others, e.g *hkb*, the enhancer activity was very transient in the ‘correct’ tissue, overlapping the gene’s expression (Supplementary Fig. 5e). The remaining five regions either had enhancer activity that did not match the interacting gene’s expression (VAChT, Olig #4), or did not match the tissue-specific interactions (Dl #2), or had no enhancer activity (Dop1R1, Toll-7 #1) (Supplementary Table S8). For example, the interacting region with the *VAChT* promoter had enhancer activity in the central nervous system, but curiously only in cells adjacent to the gene’s expression (Supplementary Fig. 5f). The Dl#2 region has enhancer activity overlapping the *Dl* gene’s expression in the endoderm and visceral muscle, but this activity does not match the predominantly neuronal-specific interaction between Dl#2 enhancer and the *Dl* promoter at 10-12h (Supplementary Fig. 5b).

These results indicate that combining developmental or tissue-specific changes in 3D promoter interactions with concordant changes in chromatin accessibility and/or H3K27ac is generally a good indicator of functional enhancer-promoter pairs. However, this is not always the case for all regions (as seen here for 26% (5/19) of tested cases), and it is not obvious why this works so well for some loci (e.g. *Oli*) and not others. Some of the interacting elements might be bystander interactions if it is a gene dense and/or very compact locus (i.e. cases where the enhancer’s activity does not match the interacting gene’s expression) or might serve a different regulatory function (i.e. cases where the element does not function as an enhancer at all). Although there was no obvious global concordance between the changes in insulator binding and the formation of differential E-P loops, differential insulator binding might contribute to facilitating some of these interactions (e.g. robo3 #1 has differential CTCF binding, Supplementary Fig. 3g).

### The nervous system has many long-range tissue-specific loops involving loci of lncRNA

Although we observed a similar propensity for tissue-specific enhancer-promoter loops in both Muscle and Neuronal tissues at 10-12h, the neuronal system often involves loops to regions overlapping (or very close to) the loci for annotated long non-coding RNAs (lncRNA). In some cases this involves promoter-promoter or enhancer-promoter loops that are only detected in neurons. For example, at the *Oli* locus, although the Oli #1 and Oli#2 interacting regions can both function as neuronal enhancers (Fig. 5b), they also both overlap the promoters of lncRNAs *CR45919* and *CR43304*, respectively. The *robo3* promoter interacts with a region (robo3 #1) overlapping *CR45006* specifically in neurons, and this region does not function as an enhancer (Supplementary Fig. 5d). Interestingly, many of these neuronal-specific lncRNA interactions are quite long-range (Supplementary Fig. 6), especially for *Drosophila*, including the *zfh1* promoter and *CR45571* (spanning 150 kb), the enhancer VT34804 and *CR46348* (spanning 164 kb), the neuronal enhancer VT10064 and *CR40341* (spanning 620 kb), and the neuronal enhancer VT58271 (adjacent to *CR45533*) and the *cut* locus (spanning 184 kb).

To determine the functional requirement of some of these newly identified neuronal specific loops for gene expression and development, we selected two for CRISPR/Cas9 deletion. In the *Oli* locus, we deleted the closest interacting element, Oli #1 (Fig. 5a, e), which includes the promoter of the lncRNA *CR45919* and two neuronal specific DHS immediately upstream of the promoter (approx. 4.2 kb). Embryos with the deleted element have no discernible effect on *Oli* expression (either the level or pattern) during embryogenesis under standard laboratory conditions, as measured by RNA *in situ* hybridization (Fig. 5e). The mutant flies are homozygous viable with no obvious locomotion defects under normal conditions. This indicates that loss of Oli #1 - one of the three interacting elements that functions as a neuronal enhancer (Oli #1-3) – is likely compensated by the other two elements (e.g. Oli #2-3), which have overlapping activity (Fig. 5b).

However, when the Oli #1 mutant embryos are raised under stress conditions (29°C), the flies have clear locomotor defects; they are unable to climb and have difficulty walking (Movie S1), two motoneuron phenotypes also observed in loss-of-function mutants for the *Olig family* gene^46^. This is therefore an example of phenotypic robustness being conferred by an apparently redundant enhancer under normal conditions, which becomes essential under stress conditions^47^. Here, we cannot discern what the role is, if any, of the lncRNAs in this locus or if they represent a form of eRNA transcribed from these enhancers.

In the *Toll-7* locus, the interacting region (Toll-7 #1) does not function as an enhancer (Fig. 5d), allowing the role of the lncRNA to be more easily assessed. We made two deletions to determine whether the proximity between the *Toll-7* promoter and the distal *CR44506* promoter region is functionally required for *Toll-7* expression and/or development. The first removed a neuronal specific DHS overlapping the *CR44506* promoter and the entire *CR44506* gene body (Fig. 5c, f, approx. 5.7 kb). The second smaller deletion removed only the *CR44506* gene body (5 kb), leaving the interacting accessible (DHS) promoter region largely intact. Both deletions had no discernible effect on *Toll-7* expression during embryogenesis under normal laboratory conditions, as seen by RNA *in situ* hybridization (Fig. 5f, shown for *CR44506-*DHS deletion), and the flies are homozygous viable and fertile. However, the majority of flies with the *CR44506*-DHS deletion are unable to fly (Movie S2), while the deletion of *CR44506* alone has a mild flight defect (Movie S3). When these mutants are raised under stress conditions (29°C), the *CR44506*-DHS mutants are unable to eclose, while all eclosing flies of the *CR44506* gene body mutant are unable to fly. In order to eclose, flies need their muscles in the thorax and legs to push open the pupal case and crawl out. Both eclosion and flightless phenotypes are commonly observed in mutants with neuromuscular defects, in keeping with the neuronal expression of the lncRNA. This indicates that the lncRNA (*CR44506*) might have a function alone, but deletion of the interacting DHS region (including the *CR44506* promoter) causes even more severe phenotypes even under normal laboratory conditions. This suggests that the promoter (or some other function of the sequence underlying the DHS) may act as a structural element to facilitate *Toll-7* regulation some 130kb away.

## DISCUSSION

Although there are many features associated with the activation of an enhancer or promoter, including transcription factor binding, chromatin state and transcription, their proximity to each other is not always correlated with their activity. For example, in early stages of *Drosophila* embryogenesis, many E-P interactions are preformed and can occur in the absence of gene activation^13^. Here, we moved to later stages of embryogenesis, examining the relationship between E-P proximity and their activity during cell fate specification and the initiation of terminal differentiation in both muscle and neurons. We selected 300 *in vivo* verified tissue-specific enhancers and 300 promoters of genes with tissue-specific expression (for either muscle or neurons), and confirmed their activity by visual inspection in the respective databases. Using this hand-curated set of *bona fide* enhancers and promoters, we compared their interaction frequencies to their activity across 5 developmental conditions using high-resolution Capture C. This enabled a comparison of interaction frequencies of enhancers and promoters within a tissue’s lineage across time (i.e. a muscle enhancer in myoblasts vs differentiated muscle), and across cell-types at the same developmental time-point (e.g. a muscle enhancer in differentiated muscle vs neurons).

During the earlier embryonic stages of tissue specification, E/P interactions are surprisingly similar between specified myoblasts and neurons, even though these cell-types have many differences in their gene expression including specific cell identity genes. These largely invariant E-P topologies are in line with our previous study, and that of others, comparing to the early blastoderm stage^13,48,49^. However, at later embryonic stages during the initiation of tissue differentiation, many new tissue-specific E/P interactions emerge, which are often more distal and are formed on top of the pre-existing landscape. Therefore, at the stages of tissue differentiation, *Drosophila* has much more dynamics in E/P contacts than previously observed at earlier stages, and is highly reminiscent to what has been observed in dissected tissue from the developing mouse embryo^16,50^.

There are several properties of these differential E/P loops that link their proximity to gene activation. First, they are tissue-specific and generally only occur in the tissue and stage in which the interacting enhancer or promoter is active. Second, they have concordance in the direction of change in E/P interaction frequencies, and in the changes of E/P chromatin state. More specifically, enhancers or promoters that go from off-to-on or on-to-off in their activity have increased or decreased interaction frequencies, DHS signal and H3K27ac levels at their interacting ‘other end’, whether it is an enhancer or a promoter. Third, even for genes that are active in both tested lineages, their underlying E/P interactions can be very different and highly tissue-specific (e.g. Delta in the mesoderm and nervous system). Fourth, of the 19 elements that we tested *in vivo*, the vast majority (74%) function as development enhancers, recapitulating part of the interacting genes’ expression.

How these dynamic tissue-specific loops are formed remains to be resolved. Although insulator binding is enriched at the interacting regions, only a fraction of the differential E/P interactions are associated with a concordant change in insulator binding. These cases usually involve a tissue-specific gain of CTCF peaks and a loss of Su(Hw) as we observed at the *Toll-7* locus, and as previously reported for *Ubx*^39^. In mammals, stable or invariant E/P interactions correlate with cohesin/CTCF binding in limb buds and activity-regulated gene expression in neuron^16,21,50^. Given the large number of insulator proteins in *Drosophila*, and therefore the large diversity in possible combinations, each locus could use a different combination of factors to control access to its regulatory elements. However, as these factors are ubiquitously expressed, they are more likely to regulate invariant E/P interactions, as proposed in the limb bud^16^. GAF/Trl, a ubiquitously expressed TF was recently reported to have such a role in flies, binding close to promoters to facilitate their E-P loops^51^.

Intriguingly, GAF/Trl and PRC1 were suggested to co-bind to tethering elements and function in their regulation. Having such a system in place could explain the pre-establishment of interactions that are invariant, and yet used for activation only in certain developmental contexts^51,52^. Our results suggest that the tissue-specific E/P interactions (which appear instructive in nature) may be regulated by tissue-specific transcription factors (TFs), as their associated DHS are enriched in the motifs for tissue-specific factors such as Mef2 in muscle-specific loops and Ttk in neuron-specific loops. This suggests a role for lineage specific TFs not only in enhancer activation, but also in facilitating E-P communication (either directly or indirectly). There is some precedence for this in other systems. Overexpression of the neuronal specific TF, Ngn2, for example, leads to increased interaction frequencies between Ngn2 bound sites (at both enhancers and their target promoter), and an increase in gene expression^26^.

In the context of embryonic progression, it is interesting to speculate why different stages of embryogenesis might require different types of E-P topologies. As development occurs very rapidly during the early blastoderm stages of embryogenesis, such pre-formed E-P topologies might facilitate very rapid gene activation, similar to what has been observed for inducible genes^19–21^. These topologies remain largely unchanged at the later stages of cell fate specification, and are remarkably similar between myoblasts and neuronal cells, even though they develop from different germ-layers. Perhaps such invariant E/P topologies facilities some plasticity in gene regulation, which is essential for trans-differentiation of cell types during embryogenesis. At even later stages, during tissue differentiation, many new tissue-specific loops are formed, which are more distal, and may require more time to become established. Genes expressed during the early blastoderm stages of development, and zygotic genome activation, tend to be short and intron less, compared to genes expressed in differentiated tissues, which can be long with complex alternative splicing. Perhaps the three-dimensional architecture of regulatory landscapes follows a similar logic, or reflects this shift in complexity. Early in embryogenesis genes rely on preformed topologies, where their enhancers and promoters act within predefined windows of interaction frequencies and tend to be more proximal. At later embryonic stages, *de novo* interactions with more distal regulatory elements add another layer to gene regulation. It is interesting to note that these are gains of new interactions, rather than pruning of the ‘older’ preexisting landscape. As E/P interactions become more cell type specific, this might facilitate robustness while at the same time help to ensure developmental lock-down during terminal tissue differentiation.

## DATA AVAILABILITY

All raw data was submitted to EMBL-EBI’s ArrayExpress (https://www.ebi.ac.uk/arrayexpress/browse.html) under accession numbers: E-MTAB-9310 (Capture C data). The processed Capture-C interaction maps, and tissue specific insulator and H3K27ac ChIP-seq peaks will be made available on the Furlong lab web page, http://furlonglab.embl.de/data.

## ACKNOWLEDGMENTS

We thank all members of the Furlong lab for discussions during the course of this project, and particularly Elena Molina, Charalampos Galouzis, Martina Varisco, and Alessandro Dulja for very useful comments on the manuscript. This work was technically supported by the EMBL Genomics Core (GeneCore), Flow Cytometry Core (FCCF) and Advanced Light Microscopy (ALMF) facilities. This work was financially supported in part by a post-doc fellowship to T.P. as part of the EI3POD programme (an EU cofounded Marie Skłodowska Curie programme (664726)), and ERC advanced grant DeCRyPT (787611), and DFG-SPP 2202 grant to E.E.F.

## AUTHOR CONTRIBUTIONS

T.P. and E.E.F. designed the study. T.P. and M.C.G. selected the enhancers/promoters, and performed the tissue-specific Capture C. T.P generated the transgenic lines, CRISPR deletions and performed nuclear DNA/RNA FISH. A.R. performed all Capture C and other data analyses with input from T.P. and E.E.F. R. M-F performed in-situ hybridizations. R.V performed the insulator and H3K27ac tissue specific ChIP-seq. A.J. designed the Capture-C probes. C.S. performed the stress analysis and movies. T.P and E.E.F. wrote the manuscript with input from all authors. All authors discussed the results and commented on the manuscript.

---

**Supplementary Figure 1:**
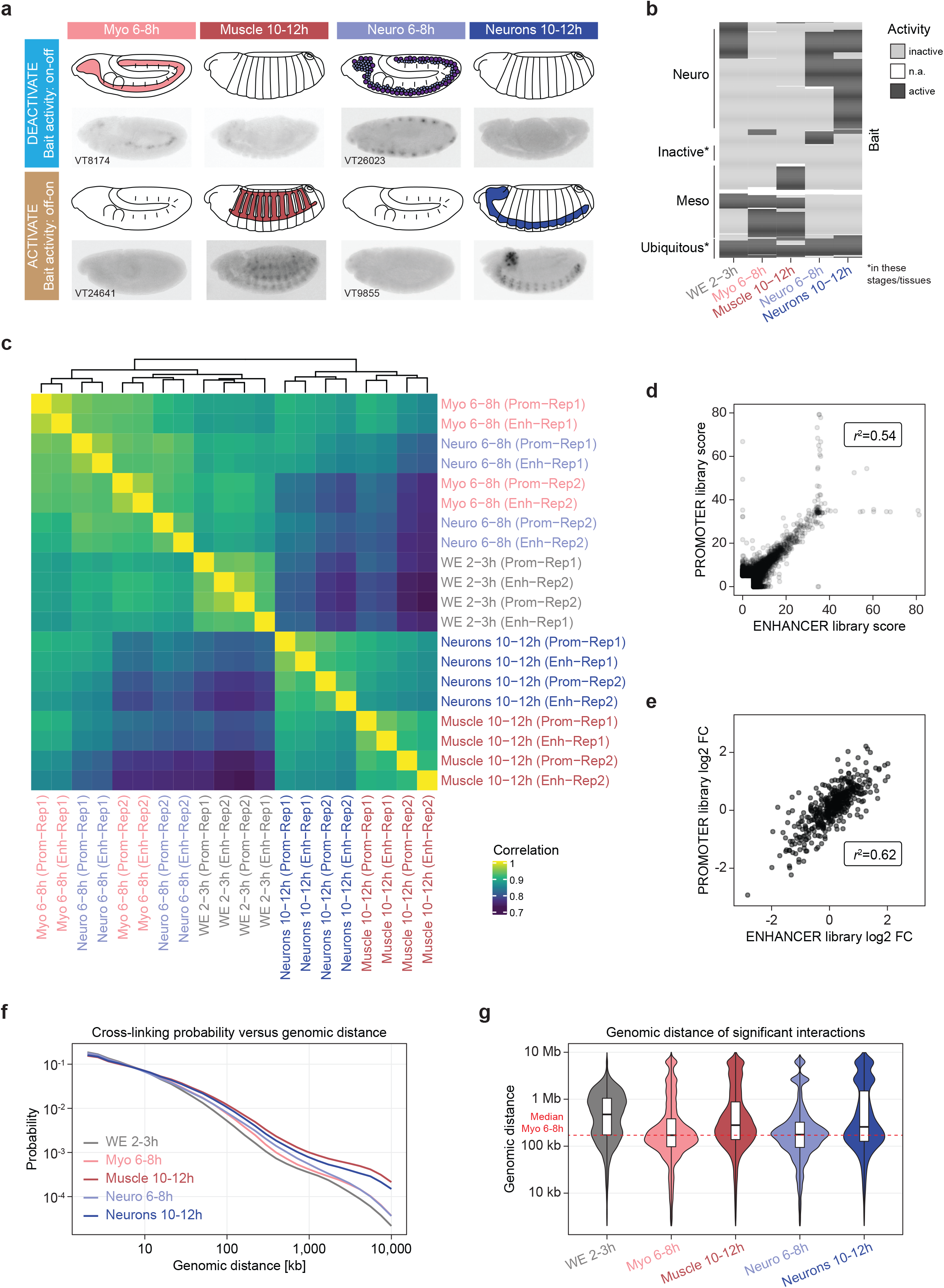
Overview of the experimental design and data quality. **(A)** Examples of developmental enhancers with dynamic tissue specific activity, which were used as baits. *Top row*: Schematic and embryo images of enhancers activate in Myoblasts (Myo) or Neuronal (Neuro) cells at 6-8h, but not at 10-12hr (on-to-off). *Bottom row*: Schematic and embryo image of enhancers not activate in Myo or Neuro at 6-8h, but are active in Muscle or Neurons at 10-12hr (off-to-on). Embryo images show *in-situ* hybridizations of the transgenic reporter, obtained from the Fly Enhancer VDRC database (https://enhancers.starklab.org/). The vast majority of enhancers and promoters were selected based on their exclusive activity in the developing mesoderm/muscle and nervous system matching one of the categories on-on, off-off, off-on, on-off. **(B)** Heatmap indicating the activity of all selected baits. Data is discretized to show active (dark grey), inactive (light grey) or unknown (n.a.) based on signal in the respective databases (e.g. VDRC, CAD4, BDGP). The respective tissue is indicated underneath. Enhancers/promoters that are dynamic in their activity between 6-8h and 10-12h (tissue specification and terminal differentiation) and tissue specific were selected. Baits were included for ubiquitously expressed or non-expressed genes as controls. Tissue-/stage-specific analysis was performed using only the baits with known and annotated activity. **(C)** Heatmap showing correlation of Capture-C interaction counts between replicates, samples, and enhancer/promoter bait libraries for each Capture-C experiment. Correlation is calculated for the 26 baits common to both bait libraries. The first level of clustering are the libraries generated from the same replicate. The data subsequently clusters by sample and then timepoint. Overall, there is very good correlation between data generated from the same sample and condition. **(D)** To assess reproducibility, we compared CHiCAGO scores generated from the 26 baits present in both the Enhancer and Promoter libraries. Overall, there is a very strong correlation between the scores generated in each library (r^2^=0.54) with clear diversion occurring only for interactions with extreme significant CHiCAGO scores (>30) in both libraries. **(E)** To assess reproducibility of the calculated log2 fold-change, we compared log2 fold-change values generated for the 26 baits present in both the Enhancer and Promoter libraries. There is a clear and strong correlation (r^2^=0.62) between the values generated in each library showing the reproducibility of both the data and its analysis. **(F)** P(s) plot displaying the probability of observing interactions at a given distance/separation between the bait and ‘otherEnd’. Over developmental time there are fewer proximal interactions (<10kb) and more distal interactions (>10kb). In the identification of differential interactions, a normalization process was applied to account for the differences in the P(s) curves. **(G)** Violin plot displaying the distribution of genomic distances between the bait and ‘otherEnd’ for all significant interactions (CHiCAGO score >5.0) identified in the 5 conditions. The dashed red line indicates the median of the distances of significant interactions in Myo 6-8h, to allow for easy comparison between conditions.

**Supplementary Figure 2:**
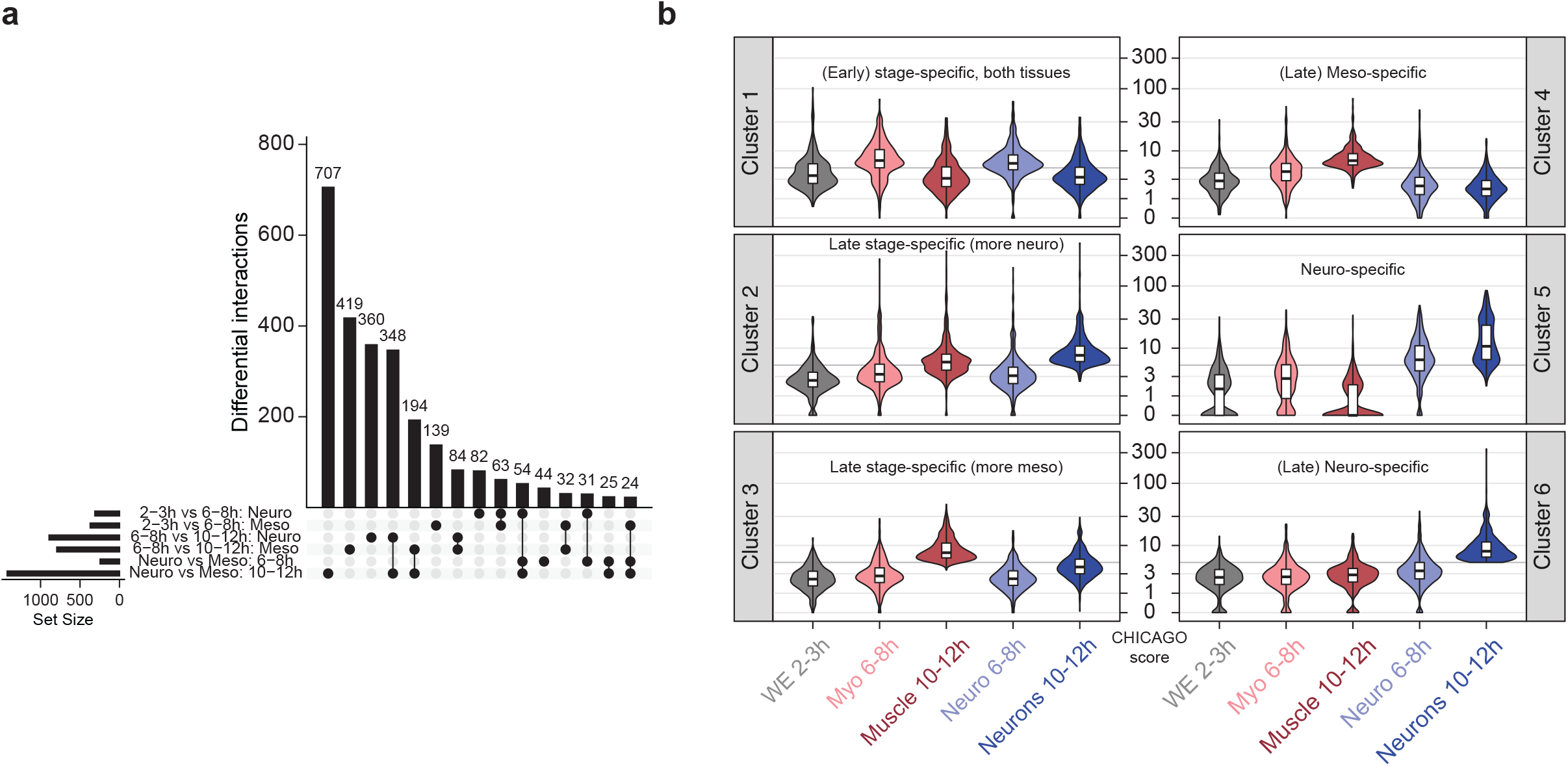
Specificity of differential interactions. **(A)** UpSet plot displaying the co-occurrence of significantly differential interactions in comparisons of different conditions. Only the 15 most frequent overlaps are shown. **(B)** Violin plots depicting the CHiCAGO scores of all interactions within a given cluster in the 5 tested conditions. The applied CHiCAGO score threshold (>= 5) is indicated by the horizontal grey line. The plot highlights that the majority of interactions in a cluster (Fig. 2D) are only significant (score >5) in the appropriate sample(s), confirming their strong enrichment in time- and/or tissue-specific interactions. In cluster 5, for example, the majority of detected interactions are only significant (score >5) in the nervous system, and have an even higher score at 10-12h. Cluster 6, the majority of interactions are exclusive to the nervous system at 10-12h (stage of differentiation).

**Supplementary Figure 3:**
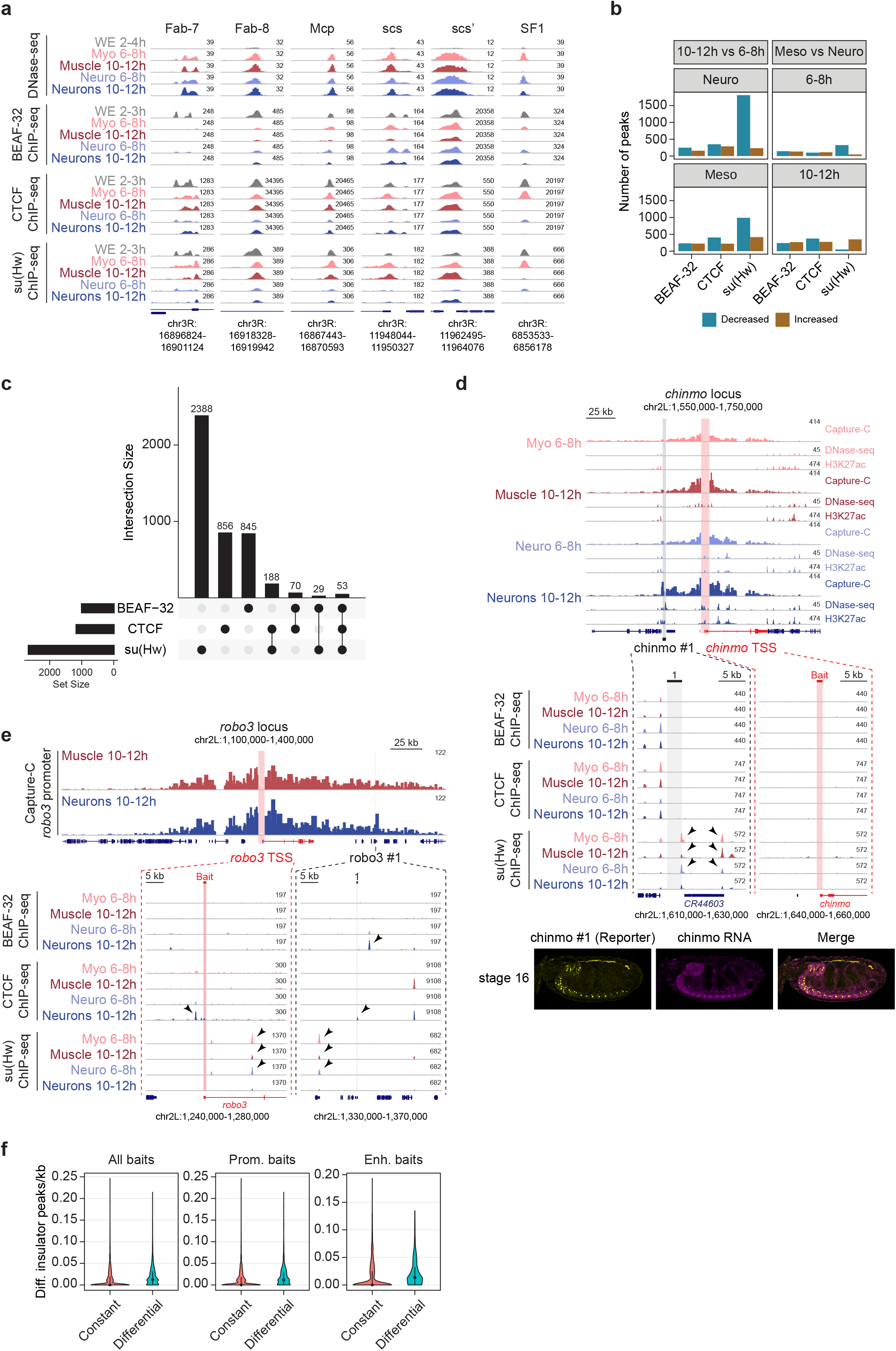
Properties of insulator binding at interacting regions. **(A)** Normalized DNase-seq and ChIP-seq signal for insulator proteins in the indicated conditions at six characterised insulator elements. Name of the element and genomic location are indicated above and below the panel, respectively. **(B)** Bar chart depicting the number of decreased (turquoise) and increased (brown) insulator peaks in the respective conditions based on differential analysis (DESeq2) of ChIP-seq signal. **(C)** UpSet plot showing the overlap of differential insulator peaks. For each insulator protein the differential peaks across all samples were merged prior to determining their overlaps. **(D, E)** Examples of differential tissue-specific interactions with tissue-specific insulator binding at the *chinmo* (**D**) and *robo3* (**E**) loci. For *chinmo*, interaction in Neuro 10-12h between the *chinmo* promoter (bait, indicated by red vertical bar) and the region around *CR44603* is associated with reduced Su(Hw) binding in Neuro 10-12h. For *robo3*, interaction between the *robo3* promoter (bait = red vertical bar) and the robo3 #1 element in Neuro 10-12h is associated with CTCF binding and reduced Su(Hw) binding (adjacent to the interacting sites) in Neuro compared to Muscle at 10-12h. **(F)** The frequency of differential insulator peaks (peaks per kb) in the interaction interval (bait to ‘otherEnd’ +/-5kb) for both constant and differential Capture-C interactions. Differential insulator peaks are found more frequently in the interaction interval of differential interactions than constant interactions.

**Supplementary Figure 4:**
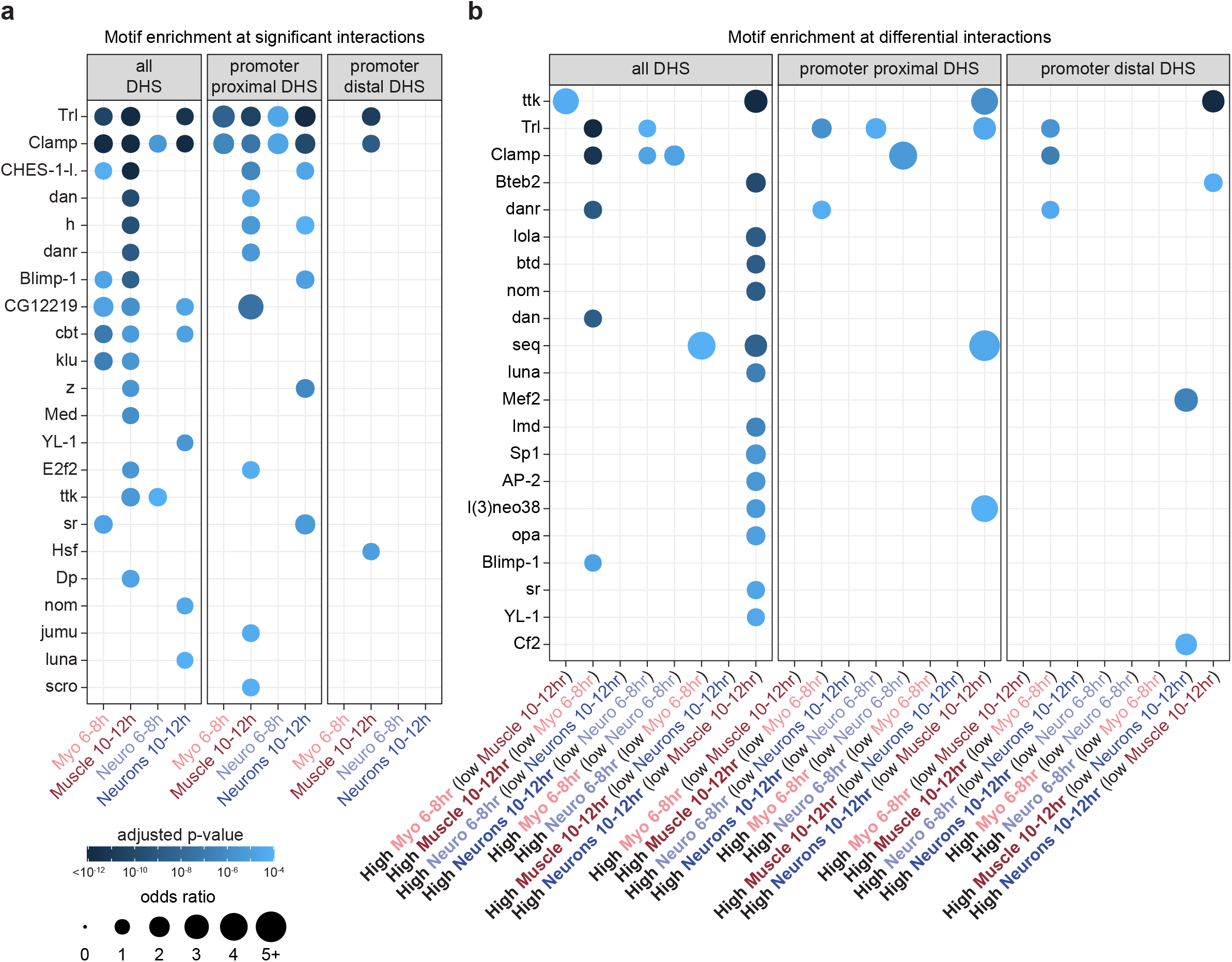
Motifs for tissue-specific transcription factors are enriched at the other ends of tissue-specific loops. **(A)** Identification of potential factors involved in the formation of E/P loops. Tissue and stage matched DHS (from Reddington et al) were divided into two groups, a test set in proximity (<500bp) to significant interactions and a control set (composed of a non-overlapping DHS set that are in proximity (<500bp) to non-significant interactions). Enrichment of *Drosophila melanogaster* motifs (from CIS-BP) in the test DHS relative to control DHS, using the AME tool. Plot shows motifs enriched in the indicated sample using an adjusted p-value cutoff of 1×10^−4^. **(B)** Motif enrichment at differential interactions. DHS were divided into three groups based on their proximity (<500bp) to increased, decreased or “other” (non-increased and non-decreased) genomic interactions characterized in the same tissue/time condition. Enrichment of *Drosophila* motifs (from CIS-BP) in either the increased or decreased DHS, relative to other DHS in the same condition, was carried out using the AME tool (doi:10.1186/1471-2105-11-165). Plots show motifs enriched in the indicated sample using an adjusted p-value cutoff of 1×10^−4^. In both (A) and (B) only DHS >10kb and <250kb from the bait were considered, and enrichments for all, promoter proximal, and promoter distal DHS (>=500bp) are shown separately.

**Supplementary Figure 5:**
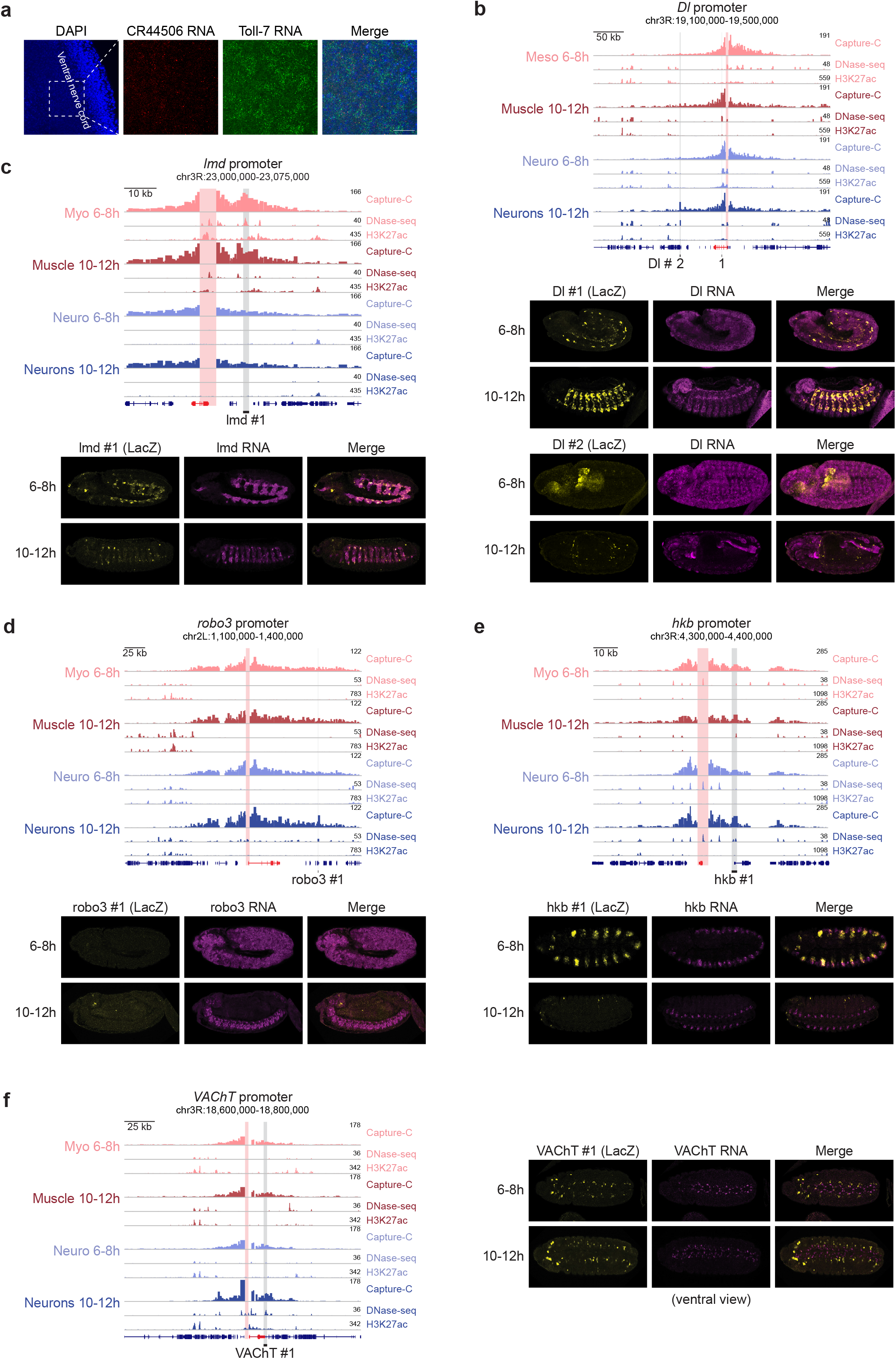
Promoter tissue-specific loops are often enhancers with activity in that tissue. **(A)** Nuclear RNA FISH (fluorescent in situ hybridization) for *CR44506* (red) and *Toll-7* RNA (green) in the ventral nerve cord of a 10-12h wildtype embryo (lateral view). The *Toll-7* signal is a mixture of nascent RNA and mRNA due to the lack of intronic sequence for hybridization to detect only nascent RNA. *CR44506* signal is faint and focal and appears to be mostly located to the nucleus at the site of transcription, and is adjacent to nuclear RNA signals for Toll-7 in some nuclei. **(B)** *Upper*: Normalized Capture-C, DNase-seq and H3K27ac ChIP-seq signal at the *Delta* (*Dl*) locus in 5 different conditions. Vertical red bar = *Delta* promoter (bait), grey bars = tested interacting region (Dl #1, Dl #2) in transgenic embryos. *Lower*: RNA *in situ* hybridization of transgenic embryos at the indicated stages for the reporter RNA (yellow, LacZ) and the Dl genes (magenta) for Dl #1 and Dl #2. **(C-F)** As in (B) for *lame duck* (*lmd*) interacting region (**C**), *roundabout 3* (*robo3*) region (**D**), *huckebein* (*hkb*) region (**E**) and *Vesicular acetylcholine transporter* (*VAChT*) region (**F**).

**Supplementary Figure 6:**
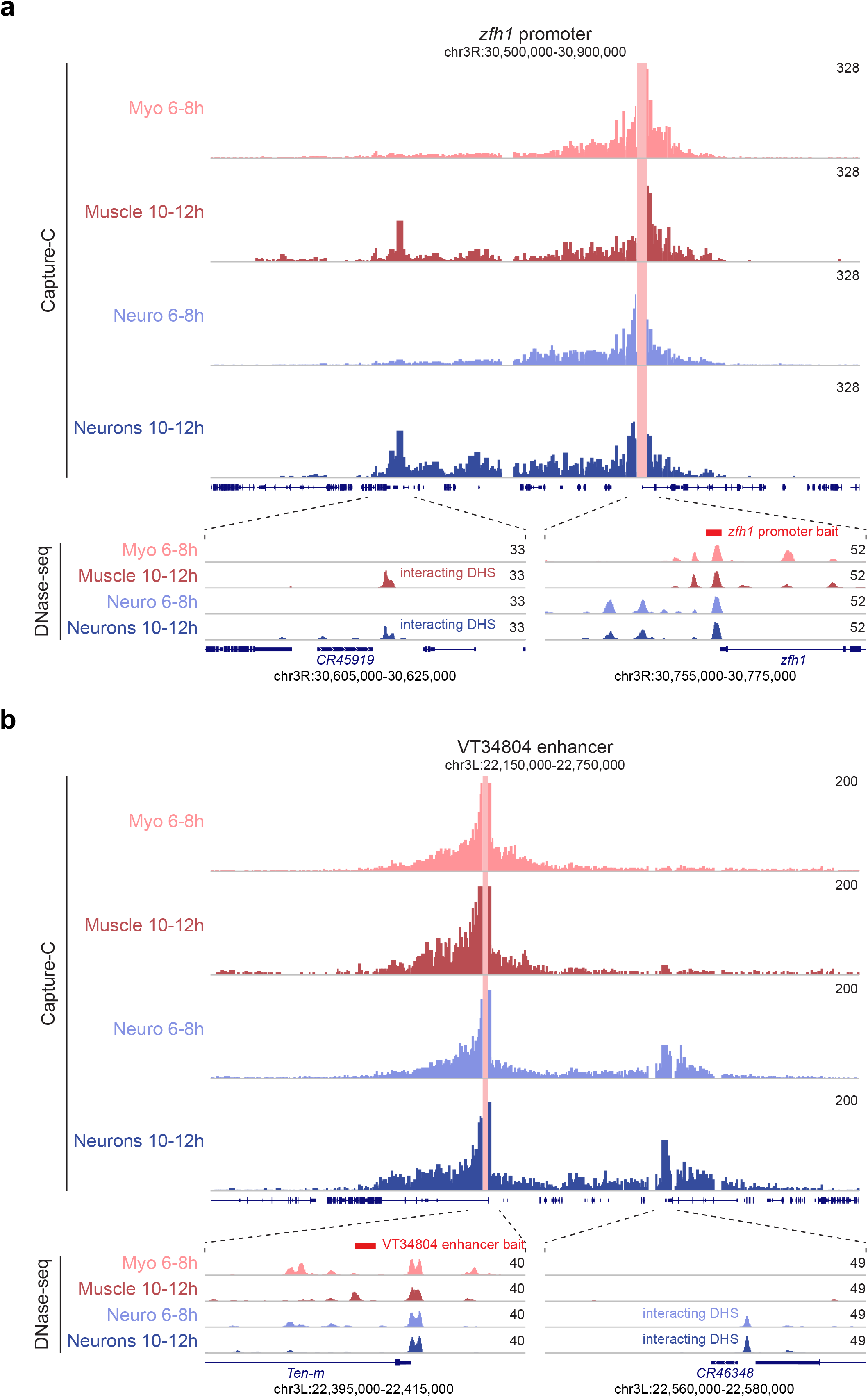
Neuronal specific E/P loops often involve loci of non-coding RNAs. **(A)** Normalized Capture-C signal at the *Zfh1* locus in 4 conditions. Vertical red bar = bait (*Zfh1* promoter). Below, zoomed in view of DHS signal in the same 4 conditions and gene models at the two ends of the loop. The lncRNA (*CR45919*) is indicated (left). **(B)** Normalized Capture-C signal from the enhancer (VT34804) used as bait (vertical red bar). Below, zoomed in view of DHS signal in the same 4 conditions and gene models at the two ends of the loop. The lncRNA (*CR46348*) is indicated (right).

## REFERENCES

1. Bharadwaj, R. et al. Conserved Higher-Order Chromatin Regulates NMDA Receptor Gene Expression and Cognition. Neuron 84, 997–1008 (2014).

2. Caputo, L. et al. The Isl1/Ldb1 Complex Orchestrates Genome-wide Chromatin Organization to Instruct Differentiation of Multipotent Cardiac Progenitors. Cell stem cell 17, 287–299 (2015).

3. Bonev, B. et al. Multiscale 3D Genome Rewiring during Mouse Neural Development. Cell 171, 557-572.e24 (2017).

4. Siersbæk, R. et al. Dynamic Rewiring of Promoter-Anchored Chromatin Loops during Adipocyte Differentiation. Molecular cell 66, 420-435.e5 (2017).

5. Madsen, J. G. S. et al. Highly interconnected enhancer communities control lineage-determining genes in human mesenchymal stem cells. Nat Genet 52, 1227–1238 (2020).

6. Mermet, J. et al. Clock-dependent chromatin topology modulates circadian transcription and behavior. Gene Dev 32, 347–358 (2018).

7. Bartman, C. R., Hsu, S. C., Hsiung, C. C.-S., Raj, A. & Blobel, G. A. Enhancer Regulation of Transcriptional Bursting Parameters Revealed by Forced Chromatin Looping. Molecular cell (2016) doi:10.1016/j.molcel.2016.03.007.

8. Deng, W. et al. Controlling long-range genomic interactions at a native locus by targeted tethering of a looping factor. Cell 149, 1233–1244 (2012).

9. Deng, W. et al. Reactivation of developmentally silenced globin genes by forced chromatin looping. Cell 158, 849–860 (2014).

10. Kim, J. H. et al. LADL: light-activated dynamic looping for endogenous gene expression control. Nature methods 16, 633–639 (2019).

11. Morgan, S. L. et al. Manipulation of nuclear architecture through CRISPR-mediated chromosomal looping. Nat Commun 8, 15993 (2017).

12. Chen, H. et al. Dynamic interplay between enhancer–promoter topology and gene activity. Nature Publishing Group 50, 1296–1303 (2018).

13. Ghavi-Helm, Y. et al. Enhancer loops appear stable during development and are associated with paused polymerase. Nature 512, 96–100 (2014).

14. Kaaij, L. J. T. et al. Enhancers reside in a unique epigenetic environment during early zebrafish development. Genome Biol 17, 146 (2016).

15. Phanstiel, D. H. et al. Static and Dynamic DNA Loops form AP-1-Bound Activation Hubs during Macrophage Development. Mol Cell 67, 1037-1048.e6 (2017).

16. Andrey, G. et al. Characterization of hundreds of regulatory landscapes in developing limbs reveals two regimes of chromatin folding. Genome Res 27, 223–233 (2017).

17. Dall’Agnese, A. et al. Transcription Factor-Directed Re-wiring of Chromatin Architecture for Somatic Cell Nuclear Reprogramming toward trans-Differentiation. Mol Cell 76, 453-472.e8 (2019).

18. Stadhouders, R. et al. Transcription factors orchestrate dynamic interplay between genome topology and gene regulation during cell reprogramming. Nature genetics 50, 238–249 (2018).

19. Jin, F. et al. A high-resolution map of the three-dimensional chromatin interactome in human cells. Nature 503, 290–294 (2013).

20. Comoglio, F. et al. Thrombopoietin signaling to chromatin elicits rapid and pervasive epigenome remodeling within poised chromatin architectures. Genome Res 28, 295–309 (2018).

21. Beagan, J. A. et al. Three-dimensional genome restructuring across timescales of activity-induced neuronal gene expression. Nat Neurosci 23, 707–717 (2020).

22. Ferri, F. et al. TRIM33 switches off Ifnb1 gene transcription during the late phase of macrophage activation. Nat Commun 6, 8900 (2015).

23. Banerjee, A. R., Kim, Y. J. & Kim, T. H. A novel virus-inducible enhancer of the interferon-β gene with tightly linked promoter and enhancer activities. Nucleic Acids Res 42, 12537–12554 (2014).

24. Javierre, B. M. et al. Lineage-Specific Genome Architecture Links Enhancers and Non-coding Disease Variants to Target Gene Promoters. Cell 167, 1369-1384.e19 (2016).

25. Qin, Y., Grimm, S. A., Roberts, J. D., Chrysovergis, K. & Wade, P. A. Alterations in promoter interaction landscape and transcriptional network underlying metabolic adaptation to diet. Nature communications 11, 1–16 (2020).

26. Noack, F. et al. Multimodal profiling of the transcriptional regulatory landscape of the developing mouse cortex identifies Neurog2 as a key epigenome remodeler. Nat Neurosci 1–14 (2022) doi:10.1038/s41593-021-01002-4.

27. Rubin, A. J. et al. Lineage-specific dynamic and pre-established enhancer–promoter contacts cooperate in terminal differentiation. Nature Publishing Group 49, 1522–1528 (2017).

28. Reddington, J. P. et al. Lineage-Resolved Enhancer and Promoter Usage during a Time Course of Embryogenesis. Dev Cell 55, 648-664.e9 (2020).

29. Kvon, E. Z. et al. Genome-scale functional characterization of Drosophila developmental enhancers in vivo. Nature 512, 91–95 (2014).

30. Rivera, J., Keränen, S. V. E., Gallo, S. M. & Halfon, M. S. REDfly: the transcriptional regulatory element database for Drosophila. Nucleic Acids Res 47, gky957. (2018).

31. Gallo, S. M., Li, L., Hu, Z. & Halfon, M. S. REDfly: a Regulatory Element Database for Drosophila. Bioinformatics 22, 381–383 (2006).

32. Gallo, S. M. et al. REDfly v3.0: toward a comprehensive database of transcriptional regulatory elements in Drosophila. Nucleic Acids Res 39, D118–D123 (2011).

33. Halfon, M. S., Gallo, S. M. & Bergman, C. M. REDfly 2.0: an integrated database of cis - regulatory modules and transcription factor binding sites in Drosophila. Nucleic Acids Res 36, D594–D598 (2008).

34. Cusanovich, D. A. et al. The cis-regulatory dynamics of embryonic development at single-cell resolution. Nature 555, 538–542 (2018).

35. Hammonds, A. S. et al. Spatial expression of transcription factors in Drosophila embryonic organ development. Genome biology 14, R140 (2013).

36. Tomancak, P. et al. Systematic determination of patterns of gene expression during Drosophila embryogenesis. Genome Biol 3, research0088.1-88.14 (2002).

37. Tomancak, P. et al. Global analysis of patterns of gene expression during Drosophilaembryogenesis. Genome Biol 8, R145 (2007).

38. Cairns, J. et al. CHiCAGO: robust detection of DNA looping interactions in Capture Hi-C data. Genome biology 17, 127 (2016).

39. Magbanua, J. P., Runneburger, E., Russell, S. & White, R. A variably occupied CTCF binding site in the ultrabithorax gene in the Drosophila bithorax complex. Molecular and cellular biology 35, 318–330 (2015).

40. Weirauch, M. T. et al. Determination and Inference of Eukaryotic Transcription Factor Sequence Specificity. Cell 158, 1431–1443 (2014).

41. Ohtsuki, S. & Levine, M. GAGA mediates the enhancer blocking activity of the eve promoter in the Drosophila embryo. Gene Dev 12, 3325–3330 (1998).

42. Schweinsberg, S. et al. The enhancer-blocking activity of the Fab-7 boundary from the Drosophila bithorax complex requires GAGA-factor-binding sites. Genetics 168, 1371–84 (2004).

43. Fuda, N. J. et al. GAGA Factor Maintains Nucleosome-Free Regions and Has a Role in RNA Polymerase II Recruitment to Promoters. Plos Genet 11, e1005108 (2015).

44. Bag, I., Dale, R. K., Palmer, C. & Lei, E. P. The zinc-finger protein CLAMP promotes gypsy chromatin insulator function in Drosophila. J Cell Sci 132, jcs226092 (2019).

45. Jordan, W. & Larschan, E. The zinc finger protein CLAMP promotes long-range chromatin interactions that mediate dosage compensation of the Drosophila male X-chromosome. Epigenet Chromatin 14, 29 (2021).

46. Oyallon, J. et al. Regulation of locomotion and motoneuron trajectory selection and targeting by the Drosophila homolog of Olig family transcription factors. Dev Biol 369, 261–276 (2012).

47. Frankel, N. et al. Phenotypic robustness conferred by apparently redundant transcriptional enhancers. Nature 466, 490–493 (2010).

48. Espinola, S. M. et al. Cis-regulatory chromatin loops arise before TADs and gene activation, and are independent of cell fate during early Drosophila development. Nat Genet 53, 477–486 (2021).

49. Ing-Simmons, E. et al. Independence of chromatin conformation and gene regulation during Drosophila dorsoventral patterning. Nat Genet 53, 487–499 (2021).

50. Chen, Z. et al. Widespread Increase in Enhancer-Promoter Interactions during Developmental Enhancer Activation in Mammals. (2022) doi:10.1101/2022.11.18.516017.

51. Batut, P. J. et al. Genome organization controls transcriptional dynamics during development. Science 375, 566–570 (2022).

52. Gindhart, J. G. & Kaufman, T. C. Identification of Polycomb and trithorax group responsive elements in the regulatory region of the Drosophila homeotic gene Sex combs reduced. Genetics 139, 797–814 (1995).

